# Systematic profiling of peptide substrate specificity in N-terminal processing by methionine aminopeptidase using mRNA display and an unnatural methionine analogue

**DOI:** 10.1101/2025.08.27.672577

**Authors:** Raphael J. Turra, Satoru Horiya, Mahesh Neralkar, Jennifer K. Bailey, Viktor Horvath, Isaac J. Krauss

**Affiliations:** Department of Chemistry, Brandeis University, 415 South St., Waltham, Massachusetts 02454-9110, United States; Wyss Institute for Biologically Inspired Engineering, Harvard University, 201 Brookline Ave., Boston MA, 02215, United States

## Abstract

Methionine aminopeptidase (MAP) is useful in chemical biology research for N-terminal processing of peptides and proteins and in medicine as a potential therapeutic target. These technologies can benefit from a precise understanding of the enzyme’s substrate specificity profiled over a wide chemical space, including not just natural substrates, peptides containing N-terminal Met, but also unnatural peptide substrates containing N-terminal Met analogues that are also cleaved by MAP like homopropargylglycine (HPG) and azidohomoalanine (AHA). A few studies have profiled substrate specificity for cleavage of N-terminal Met, but none have systematically done so using N-terminal Met analogues. Therefore, we devised a high-throughput profiling experiment based on mRNA display and NGS to probe MAP’s substrate specificity using N-terminal HPG. From subgroup analysis of either single residues or two-residue combinations, we could establish the impact of residue identity at various positions downstream from the cleavage site. To validate the selection results, a collection of short peptides was chemically synthesized and assayed for cleavage efficiency, where we observed reasonable agreement with selection data. Results generally followed previously reported trends using N-terminal Met, the strongest trend being that the second residue (P1’ position) had the greatest impact on MAP cleavage efficiency with moderate impacts discerned for residues further downstream which could be rationalized through modeling the enzyme-substrate interaction.

## Introduction

Methionine aminopeptidase (MAP) has demonstrated utility as both a tool for chemical biology as well as a potential therapeutic target. The natural function of MAP is to cleave the obligatory N-terminal Met residue from nascent peptides either alone (as in eukaryotes) or in conjunction with peptide deformylase (PDF) (as in prokaryotes) (**Fig. 1a**). Medically, inhibition of bacterial MAP is a potential antibiotic strategy, while inhibition of human MAP2 may be beneficial for combatting obesity and cancer.^1,2^ In chemical biology research, noncanonical amino acid (ncAA) Met analogues like homopropargylglycine (HPG) and azidohomoalanine (AHA) have been used to residue-specifically incorporate chemical handles into proteins for bioorthogonal coupling of probes and other functional conjugates.^3,4^ Since the activity of MAP extends to cleavage of N-terminal Met analogues, MAP enables removal of an N-terminal conjugation site when only internal conjugation is desirable (**Fig. 1b**). For example, we recently employed MAP in the construction of glycopeptide mRNA display libraries in which CuAAC glycan attachment occurred only at internal sites.^5^

**Figure 1:**
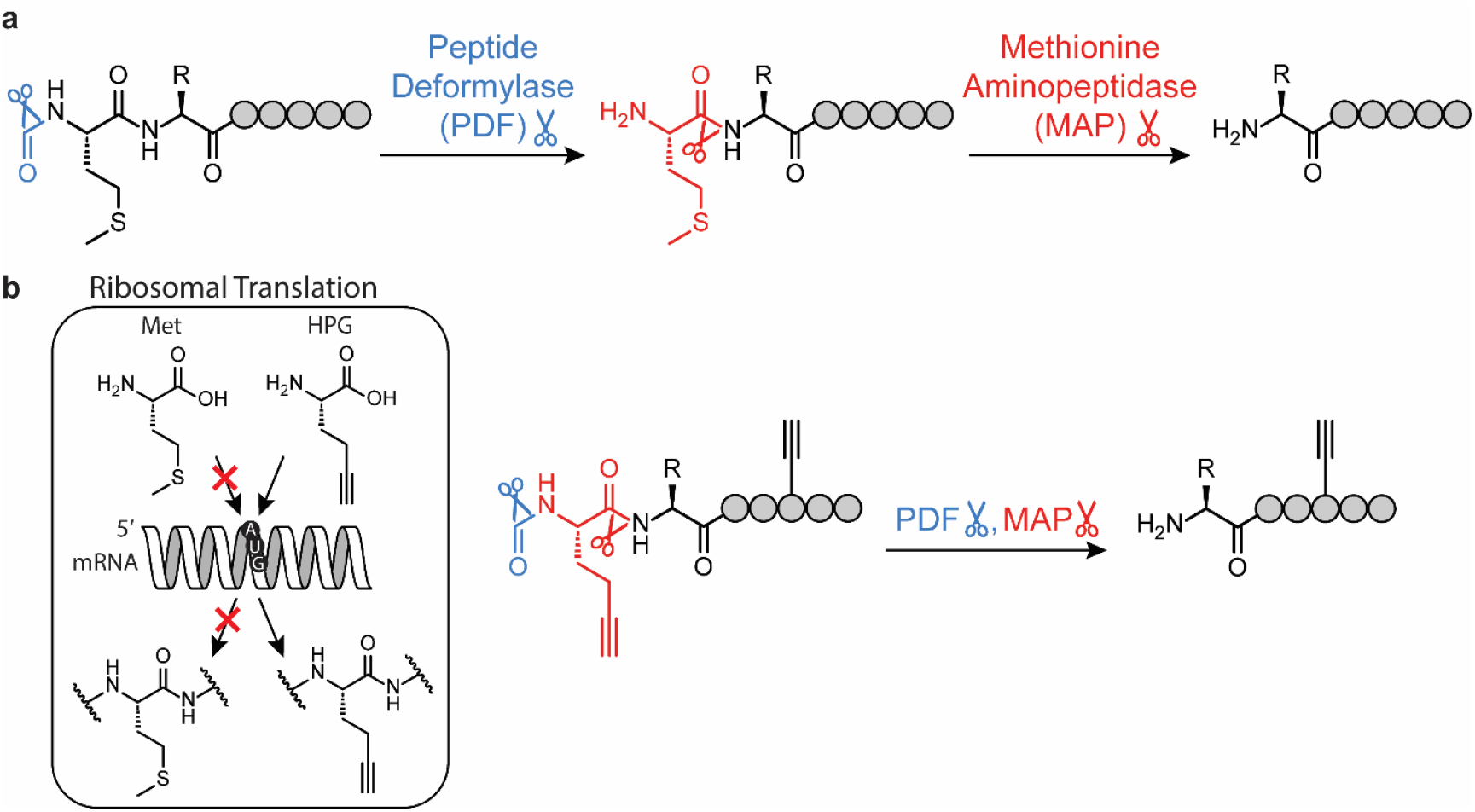
(a) N-terminal processing in prokaryotes by the combined activities of PDF and MAP to cleave formyl-Met. (b) Analogous N-terminal processing occurs with methionine analogues like formyl-HPG in the case of residue-specific noncanonical amino acid incorporation.

Underlying all these applications is the need to understand MAP’s substrate preference to ensure either complete cleavage or complete inhibition of cleavage. Previous work on this question has focused on MAP’s natural substrates, peptides containing N-terminal Met. The most comprehensive of these studies confirm that while the identity of the residue at the second position (P1’) has the greatest impact on cleavage success, the identity of residues further downstream can also have a notable impact.^6,7^ While N-terminal Met has been thoroughly studied,^6– 10^ no such comprehensive and systematic study has been conducted to determine MAP substrate specificity of peptides containing N-terminal Met analogues. Smaller studies so far have demonstrated that while the rules determining cleavage efficiency appear similar to those for peptides containing N-terminal Met, there may be some critical differences.^11–13^

We opted to use mRNA display to profile the substrate specificity of MAP. In general, the combinatorial and high-throughput nature of display techniques allow for interrogation of a large chemical space, and several studies have demonstrated that display techniques in conjunction with next-generation sequencing (NGS) can elucidate the substrate preference of promiscuous post-translational enzymes.^14–18^ As an *in vitro* technique, mRNA display has the advantages of extreme flexibility regarding noncanonical amino acid incorporation and post-translational (including chemical) modifications as well as superior diversity unrestrained by transformation efficiency allowing for more complete sampling of a wider chemical space.^19^ Moreover, our use of PURE system (Protein synthesis Using Recombinant Elements) enables facile incorporation of unnatural amino acids^20–22^ as well as control over the presence/absence of N-terminal modification enzymes in translation reactions. In this work, we present the results of an mRNA display selection designed to select for sequences based on their differential ability to undergo MAP cleavage of their N-terminal HPG. Briefly, short peptide-encoding mRNA display libraries containing an N-terminal HPG were subjected to CuAAC click attachment of a biotinylated azide linker and partitioned using streptavidin magnetic beads. Partitioned fractions were sequenced, and relative abundance in sequencing results was used to calculate the extent of HPG cleavage for comparison of different library subgroups.

## Results and Discussion

### Library Design

For the profiling experiment, four short peptide-encoding DNA libraries were prepared, each containing a randomized region consisting of NNS codons (N = A/T/G/C, S = G/C) (**Fig. 2**). The N-terminal HPG (*M*) encoded by each library was followed by either Ala or a random amino acid in the P1’ position (“Fixed” and “Variable” libraries, respectively) and then either 8 or 4 random amino acids (“Long” and “Short” libraries, respectively), followed by a linker and His tag. Short libraries were limited in length to ensure complete coverage by NGS whereas Long libraries were extended to test further downstream positions. All libraries began with a constant 5’ region containing a T7 promoter, epsilon enhancer, and Shine-Dalgarno sequence and ended with a constant 3’ sequence for crosslinking a puromycin-containing oligonucleotide. To prevent library cross-contamination, Variable and Fixed libraries were designed with different 3’ constant regions such that they used different PCR and reverse transcription primers (**Supplementary Table S1**).

**Figure 2:**
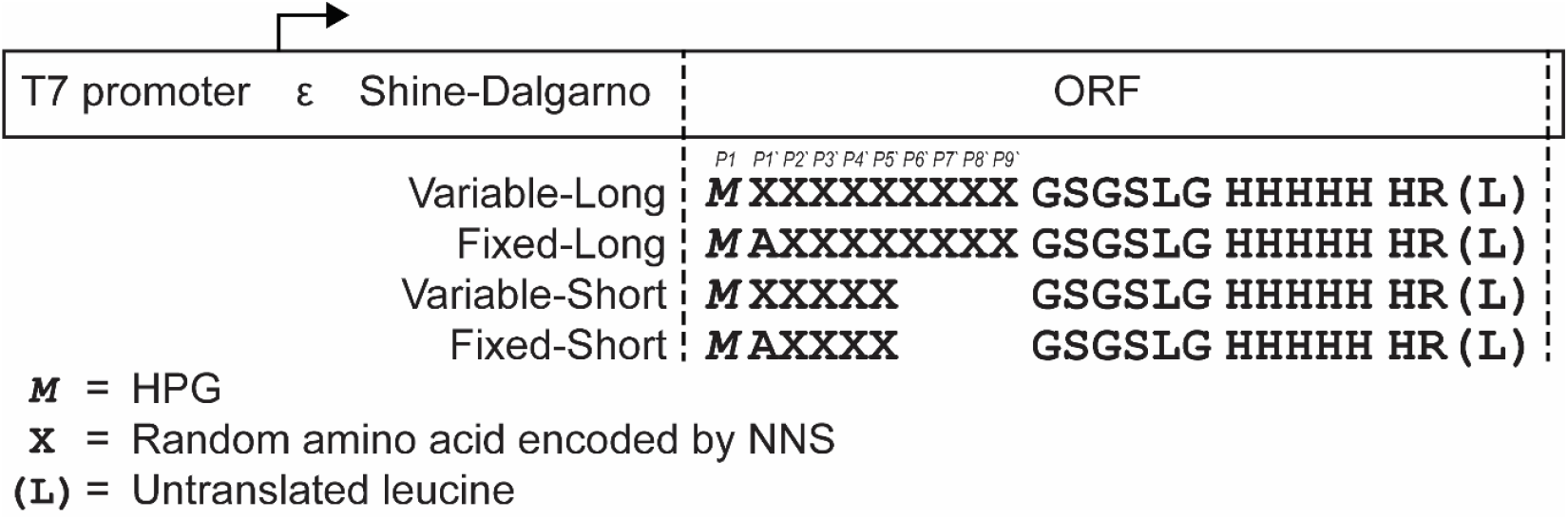
MAP library design. The final leucine codon is not translated as it is the site of photochemical attachment of the puromycin oligo for mRNA display.

### mRNA Display Selection

Overall, the profiling experiment consisted of two rounds of selection in which libraries were biotinylated by CuAAC and retrieved on streptavidin beads; the first round was performed without MAP digestion to enrich for “clickable” sequences that could be successfully biotinylated by CuAAC, and the second was performed with MAP digestion to partition cleavable from uncleavable sequences (**Fig. 3**). In the second round, libraries were split, and a parallel control selection was performed without MAP digestion to control for differential click efficiency of various sequences.

**Figure 3:**
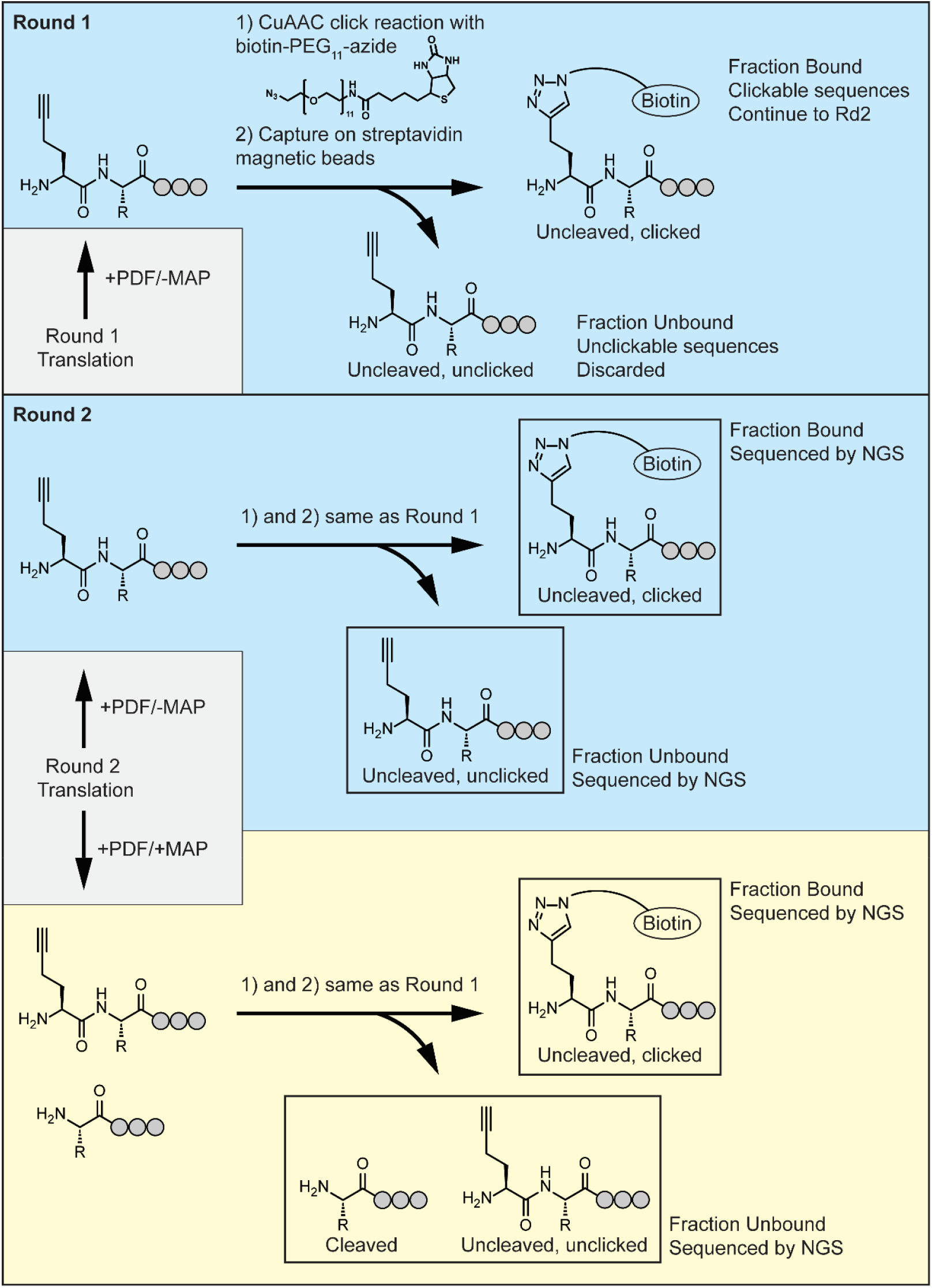
Selection scheme. Library fractions sequenced by NGS are indicated by boxes.

For Round 1 of selection, library preparation largely followed the procedure used in our previous studies.^5,23^ DNA libraries were transcribed by T7 RNA polymerase, and the resulting RNA libraries were photo-crosslinked to a puromycin-containing oligo. Purified crosslinked RNA libraries were translated with PURE system containing Met analogue HPG in the place of Met, generating mRNA-alkynyl peptide fusions. Fusions were captured on streptavidin Ultralink™ resin loaded with photocleavable biotinylated reverse transcription primers, beads were separated from unfused peptides, and reverse transcription and digestion with PDF were performed on-resin (**Supplementary Fig. S1**). Although we were previously able to N-terminally process long peptide libraries by inclusion of MAP and PDF in the translation mixture,^5^ in this case, it was necessary to perform N-terminal processing as a post-translational enzymatic reaction (**Supplementary Fig. S2**). We hypothesize that this is because these peptides were too short to reach the end of the ribosomal exit tunnel,^24,25^ making their N-terminus inaccessible for cleavage. Fusions were recovered from the resin after photocleavage of the linker with 365 nm UV light. After Ni-NTA purification to remove any unfused RNA and DNA, fusion libraries were subjected to a negative selection against M280 streptavidin magnetic beads to remove peptide sequences that bind to the beads. Remaining library was subjected to CuAAC to attach a biotin-PEG_11_-azide linker.

In Round 1 of selection (without MAP cleavage), we noticed that fraction bound values, which would be 100% in the case of perfect click efficiency and bead uptake, were consistently between 38 and 53% (**Supplementary Table S2**). Optimization experiments consisting of resubjecting libraries to a second round of click reaction before streptavidin pulldown or trying different streptavidin resins did not improve fraction bound values, so selection was carried forward as is.

For Round 2 of selection, DNA was recovered from the Round 1 bound fraction by on-bead PCR, and cDNA-mRNA-peptide fusion libraries were prepared in the same way as in Round 1 with a few differences. Libraries were split in half after translation for a total of 8 libraries prepared in parallel. While the first half was subjected to just PDF digestion while on streptavidin Ultralink™ resin, the second half was subjected to both PDF and MAP digestion. Some MAP-digested libraries demonstrated unusually high binding during negative selection against M280 streptavidin magnetic beads (up to 11% fraction bound), so a second negative selection with streptavidin beads was performed, in which we observed minimal binding in all libraries. After verifying that libraries could be successfully functionalized on a test scale with a large oligosaccharide azide previously used by the lab (**Supplementary Fig. S3**), libraries were then twice subjected to CuAAC with biotin-PEG_11_-azide and incubated with M280 streptavidin magnetic beads. Non-MAP-digested libraries demonstrated fraction bound values from 45 to 53%, very similar to those observed for Round 1, while MAP-digested libraries demonstrated lower fraction bound values from 5 to 27% (**Supplementary Table S2**), consistent with the expected removal of the N-terminal HPG alkynyl click partner. As predicted, Fixed libraries demonstrated much lower fraction bound values than Variable libraries likely owing to the fixed Ala in the P1’ position known to be advantageous for MAP cleavage. DNA was recovered from both bound and unbound fractions and sequenced by NGS (Illumina HiSeq, GENEWIZ).

### NGS Analysis

NGS results were processed to identify all unique peptide sequences in each library and collect their frequencies. The 16 library DNA samples (4 libraries × +/-MAP × bound/unbound fractions) were labeled with different adapter barcodes and pooled for sequencing. Starting from the raw Illumina paired-end reads reported in FASTQ files, we used Paired-End reAd MergeR (PEAR v0.9.11)^26^ to assemble the reads using a PHRED quality score cutoff of 36 for trimming, p-value of 0.001, and minimum overlap of 100 bp. Typically, 80-91% of reads were successfully assembled. Integrating SeqKit (v.2.2.0)^27^ for FASTQ/FASTA file manipulation, a collection of AWK and BASH scripts were used to orient the fragments into the forward direction, crop off the 5’ and 3’ constant regions, and align the sequences at their start codon. Nucleotide sequences were then translated into peptide sequences, and those sequences containing any combination of HPG or stop codon in the randomized region, amino acid other than HPG in the P1 position, or “X” (indicating the original nucleotide sequence included an uncalled base) were filtered out. While the proportion of HPG in the randomized region was highly variable across libraries, the combination of sequences filtered out based on other conditions never exceeded 3.5%.

NGS data could accurately quantify the proportions of different subgroups in each library, but two types of corrections needed to be made in the analysis. First, because sequences in each library fraction compete for sequencing, we used scintillation counting to quantify the total amount of peptides (*n*_*x*_) in each fraction (*x* = {U, B, dU, dB} where U = Unbound, B = Bound, dU = MAP-digested Unbound, dB = MAP-digested Bound) as given in **Supplementary Table S3**. For each peptide we expected one cDNA molecule displayed and uniformly PCR amplified across the library for sequencing such that proportionality exists between number of reads (*N*) and amount of peptides (*n*) as defined in equation (1). The peptide amounts of each subgroup (*n*_*x,i*_, where *i* denotes amino acid residue at position P1’ or P2’, etc.) is a fraction of the total amount of peptides (*n*_*x*_) proportional to the ratio between the number of reads of a subgroup (*N*_*x,i*_) and the total number of reads (*N*_*x*_) of a library as defined in equation (2). (1)

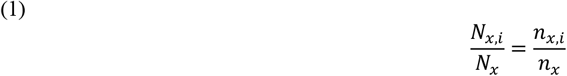

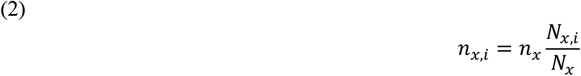

Second, due to the observed variability in the click efficiency and effectiveness of the pull-down procedures, the molar amounts of cleaved and uncleaved peptides were not treated as simply proportional to their NGS frequencies. To correct for differential efficiency of click biotinylation and/or pulldown, we defined subgroup-specific click efficiency factors (*f*) calculated from bound versus unbound peptide quantities in Round 2 non-MAP-digested libraries as shown in equation (3).

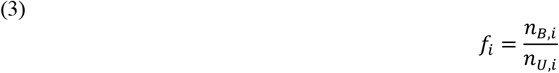

The amount of peptides of a subgroup that were bound or not bound to the beads can be calculated using equation 2. The bound amount *(n*_*dB,i*_) corresponds to the peptides that the MAP enzyme failed to cleave and was successfully clicked, and the unbound amount (*n*_*dU,i*_) corresponds to the peptides that were cleaved plus those that were cleaved but failed to click. By utilizing the click efficiency factor (eq. 3) we can estimate the cleaved but unclicked portion of the unbound peptides, which then can be used to find the total amount of cleaved peptides 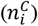 in a subgroup as shown in equation 4.

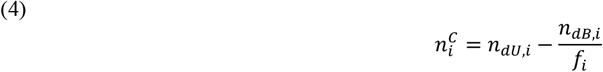

Similarly, the total amount of peptides that remained uncleaved can be found using equation 5. (5)

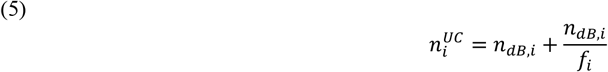

We define cleavage efficiency (*E*) in a subgroup as the cleaved portion of peptides of the total peptide amount where click chemistry was expected to be successful as defined in equation (6).

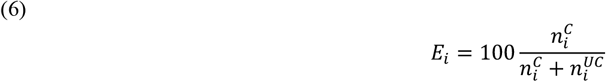

We first examined the impact of the P1’ position on MAP cleavage efficiency. Since Fixed libraries contained a fixed Ala in the P1’ position, percent cleaved calculations were only conducted on the Variable-Long and Variable-Short libraries. Sorting groups with each P1’ amino acid by percent cleavage, three tiers are evident (**Fig. 4a**). The first tier, made up of P1’ = Ser, Pro, Thr, Ala, Gly, and Val, reaches the highest level of cleavage, approximately 85% across all sequences. The second tier, made up of P1’ = Asn, Ile, Gln, Cys, Glu, and Leu, exhibit intermediate (40-75%) levels of cleavage. The third tier, made up of P1’ = Asp, His, Phe, Trp, Lys, Tyr, and Arg, are cleaved very little (≤ 30%). Results from the Long and Short libraries were highly correlated, highlighting the consistency of the profiling experiment (**Supplementary Fig. S4**). It should be noted that, based on previous literature, Tier 1 and Tier 3 subgroups were expected to cleave completely or not at all, while we actually observed values of 85% and 20% cleavage, respectively. This discrepancy may result from the fact that these are average cleavage values of diverse peptides within each subset; for instance, sequences with P1’ = Ala should normally be easily cleaved, but we found they were not if they contained Cys at P2’ or P3’. Additionally, biases could result from imperfect correction of noise introduced by click efficiency, PDF digestion bias, on-resin N-terminal processing, differential bead uptake efficiency, or other biases inherent to mRNA display selections. Regardless of the absolute scaling of the values, numerous trends based on relative ranking are clear and can be structurally rationalized.

**Figure 4:**
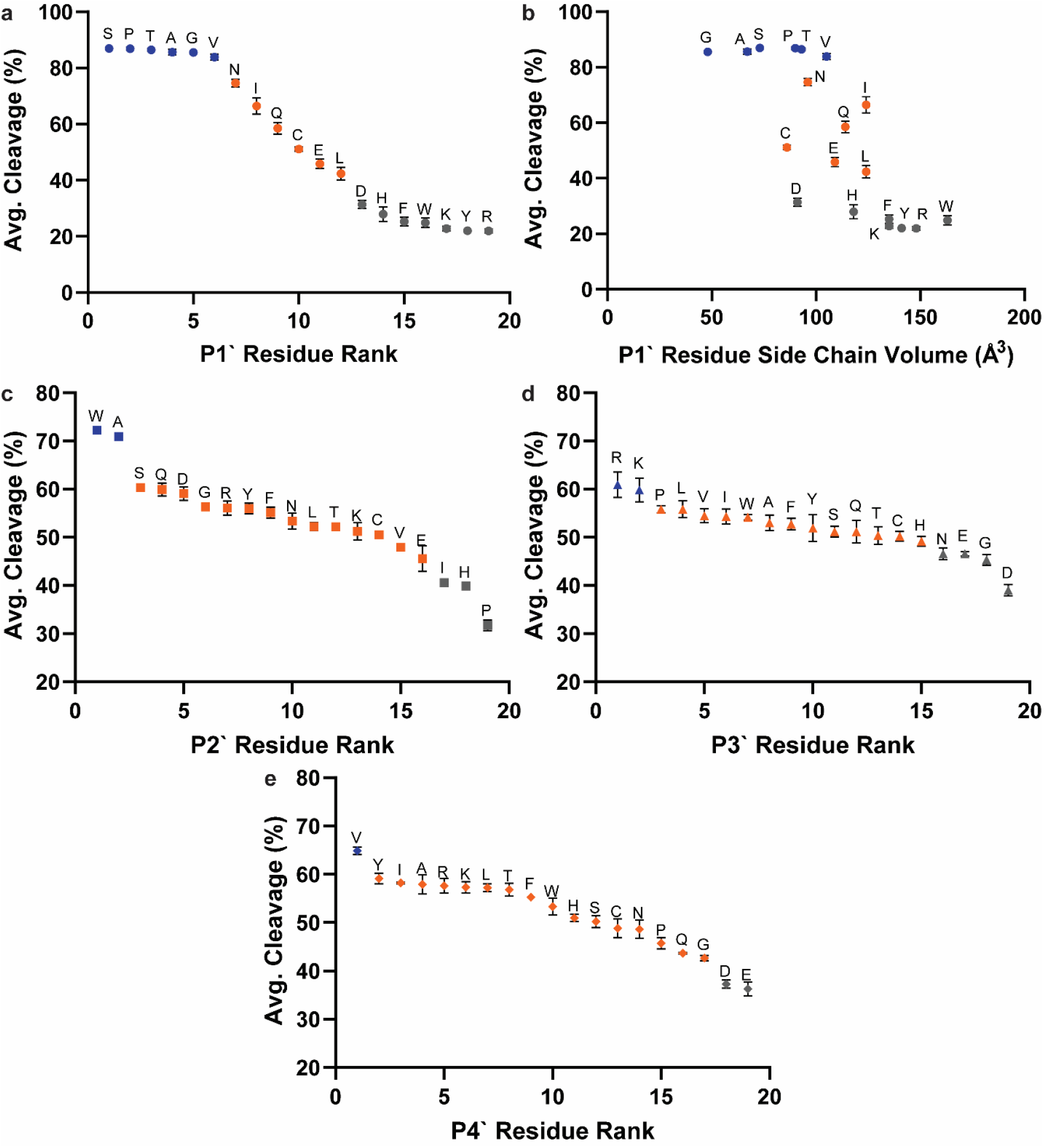
Estimated percent cleavage values from subgroup analyses. Values from P1’ subgroup analysis are presented as a ranked list (a) and plotted against residue side chain size (b). Values from P2’ (c), P3’ (d), and P4’ (e) subgroup analyses with selected P1’ = {Asn, Ile, Gln, Glu, Leu, or Asp} are presented only as ranked lists. Colors are used to qualitatively define cleavage efficiency tiers, with blue assigned to Tier 1, orange to Tier 2, and gray to Tier 3. Data are averaged between Variable-Long and Variable-Short libraries, and error bars represent the standard deviation. Individual library data are given in **Supplementary Figs. S4**,**6-8**.

Among the three tiers noted above, side chain volume has a strong impact on cleavage efficiency. Plotting amino acid percent cleavage against side chain volume reveals a clear sigmoidal trendline reaching a high plateau at small side chain size and a low plateau at large side chain size (**Fig. 4b**). Looking at side chain structures, each addition to the side chain length is detrimental to cleavage efficiency (*i*.*e*. Val vs Leu/Ile, Asn vs Gln). Compared with alkyl side chains of similar size, a polar but uncharged moiety appears to improve cleavage efficiency (*i*.*e*. Val vs Thr, Leu vs Asn), potentially by allowing for additional stabilizing H-bonds. Acidic P1’ side chains are detrimental to cleavage efficiency (*i*.*e*. Gln vs Glu, Asn vs Asp), with Asp being worse for cleavage than Glu despite having a shorter side chain. Aromatic and basic amino acids at P1’, which all have long or large side chains, all effectively inhibit MAP cleavage in the P1’ position.

It is interesting to note that, excluding Cys, the click efficiency factor values for each P1’ amino acid lay within a narrow range: *f* = 0.65 to 0.85 and 0.95 to 1.15 for the Variable-Long and Variable-Short libraries, respectively. This suggests that click efficiency was more influenced by library length than by the identity of the P1’ amino acid, excluding Cys, which was especially detrimental to click efficiency (*f* = 0.36 and 0.51 for Variable-Long and Variable-Short libraries, respectively). Among the other P1’ amino acids, despite the relatively small range, *f* values were nonetheless significantly correlated between Long and Short libraries (**Supplementary Fig. S5**).

Subgroup analyses were also conducted for P2’ and later positions in both Fixed and Variable libraries, but observed impacts on cleavage efficiency and correlations between libraries declined as distance from the cleavage site increased. As would be expected from previous observations, the percent cleavage values were distributed over a smaller range for the P2’ position, reflecting the large impact of the P1’ position, which is averaged in this type of subgroup analysis. To highlight the effect of the P2’ and downstream positions, we focused our analysis on peptides containing P1’ with an intermediate effect on cleavage (Asn, Ile, Gln, Glu, Leu, or Asp). From the sorted percent cleavage graph of the P2’ position, the best two (Trp and Ala) and worst three (Pro, His, and Ile) P2’ amino acids for MAP cleavage are evident (**Fig. 4c**), with high agreement maintained between Variable-Long and Variable-Short libraries (**Supplementary Fig. S6c**). Interestingly, the two best amino acids in the P3’ position are basic (Arg and Lys) while two of the worst amino acids are acidic (Glu and Asp), along with Gly and Asn (**Fig 4d, correlation in Supplementary Fig. S7c**). In the P4’ position, acidic residues (Glu and Asp) were again disfavored whereas Val appeared enriched (**Fig. 4e, correlation in Supplementary Fig. S8c**). Consistent patterns beyond the P4’ position were difficult to discern because of even smaller ranges in cleavage efficiency and increases in variation between Variable-Long and Variable-Short libraries. Except for Cys in most positions, we observed high cleavage efficiencies in all Fixed library subgroups, so we could not discern clear patterns beyond the fixed P1’ position (**Supplementary Fig. S9**). Cys was a consistent outlier in Fixed library analyses, resulting in lower cleavage efficiency when present at any position.

To help rationalize some of these observations, we examined models of the enzyme interaction with various peptide substrates. Starting with a published crystal structure of MAP (PDB: 2MAT)^28^, the electrostatic potential surface as calculated by Adaptive Poisson-Boltzmann Solver (APBS)^29,30^ revealed negatively charged surface at the periphery of the binding pocket (**Fig. 5a**). We then used AlphaFold 3 through AlphaFold Server^31^ to model enzyme-substrate interactions with a collection of hexapeptides. The complex with what should be a highly efficiently cleaved hexapeptide, MAWRVS, shown in **Fig. 5b**, was modeled with high confidence (ipTM = 0.96, pTM = 0.97) and is consistent with several of the substrate preferences observed in the selection. Met fits into the confirmed binding pocket observed in several crystal structures,^32,33^ and the P1’ side chain has very little space, consistent with the preference for small P1’ residues. Interestingly, the P2’ Trp side chain efficiently fills a groove on the enzyme surface, consistent with the high cleavage activity of P2’ Trp. Since the positively charged P3’ Arg is oriented along the negatively charged surface of the enzyme, this structure suggests electrostatic attraction as a factor favoring Arg and Lys at this position. Conversely, electrostatic repulsion of negative sidechains in P3’ Glu and Asp is consistent with the lower cleavage of these substrates. Among Glu and Asp, the longer side chain of Glu is better able to distance itself from negatively charged binding pocket and is less detrimental to cleavage (**Supplementary Fig. S10**). Electrostatic repulsion is also consistent with the slower cleavage of substrates with negative P4’ residues, although this is less obvious from the structure.

**Figure 5:**
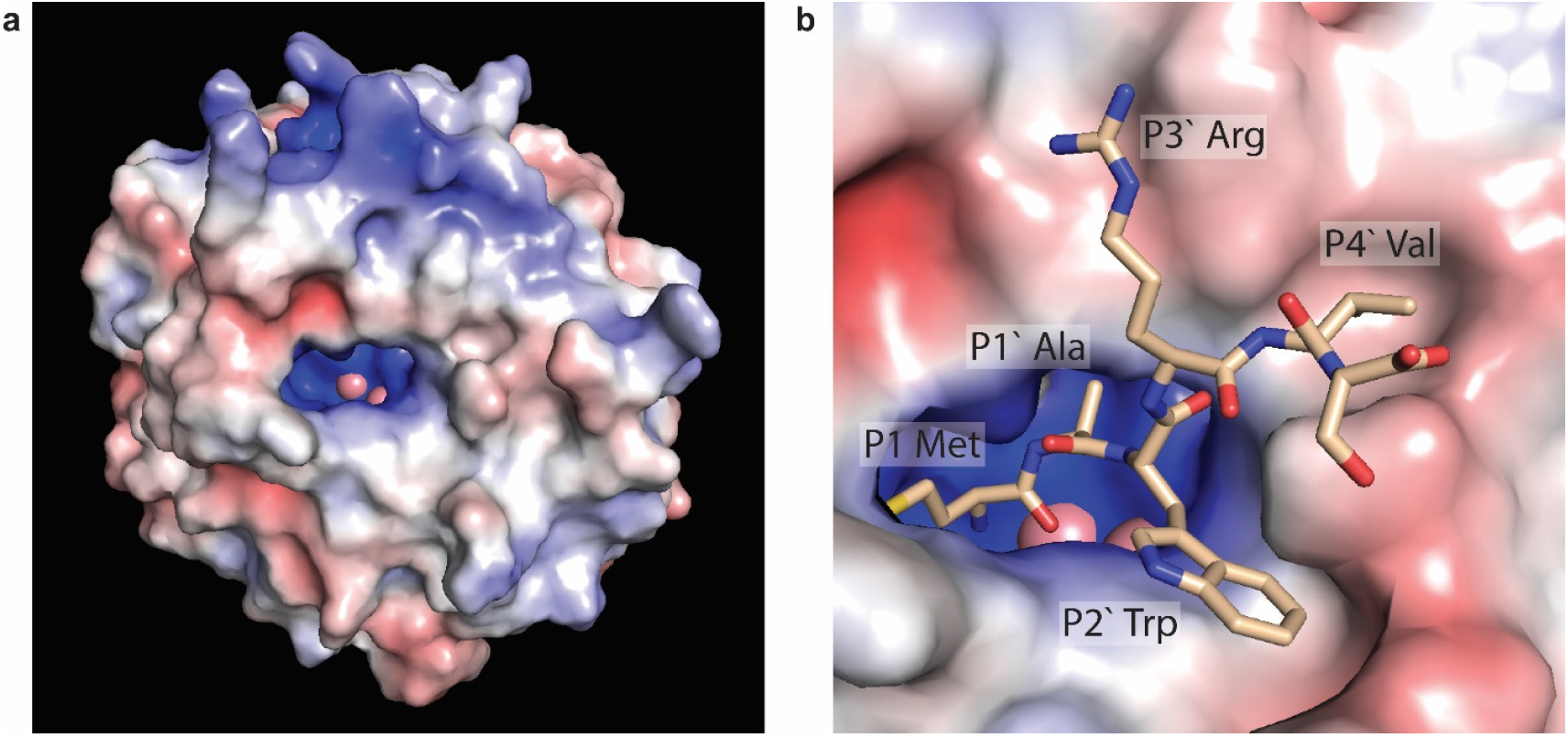
(a) Electrostatic potential surface of the MAP enzyme as calculated from published crystal structure (PDB: 2MAT)^28^ using PyMOL 1.8 with APBS plugin^29,30^. (b) AlphaFold predicted structure (ipTM = 0.96, pTM = 0.97) of MAP (colored by electrostatic potential calculation) in complex with short peptide MAWRVS (see **Supplementary Fig. S10** for other peptides modeled).

Since the amino acid residue at the P1’ position has the strongest impact on cleavage efficiency, further subgroup analyses split libraries into various P1’ × P2’ combinations or P2’ × P3’ combinations with a constant P1’ position. For the P1’ × P2’ analysis, percent cleaved for each combination was well correlated between Variable-Short and Variable-Long libraries (**Supplementary Fig. S11a**) and plotted on a heatmap (**Fig. 6a**) sorted by average values across columns and rows. As expected from the previous P1’ subgroup analysis, Tier 1 amino acids and Tier 3 amino acids demonstrate universally high and low cleavage efficiencies, respectively, regardless of the amino acid in the P2’ position. From the heatmap we can also observe that Ala and Trp in the P2’ position promote cleavage while Pro in the P2’ position suppresses it. These trends are primarily visible when looking at the impact these P2’ amino acids have on P1’ Tier 2 amino acids in the center columns of the heatmap. Cys behaves differently from other amino acids with a consistently intermediate cleavage efficiency when in the P1’ position and, when in the P2’ position, inhibitory impact on sequences with P1’ Tier 1 amino acids but promoting impact on sequences with P1’ Tier 3 amino acids.

**Figure 6:**
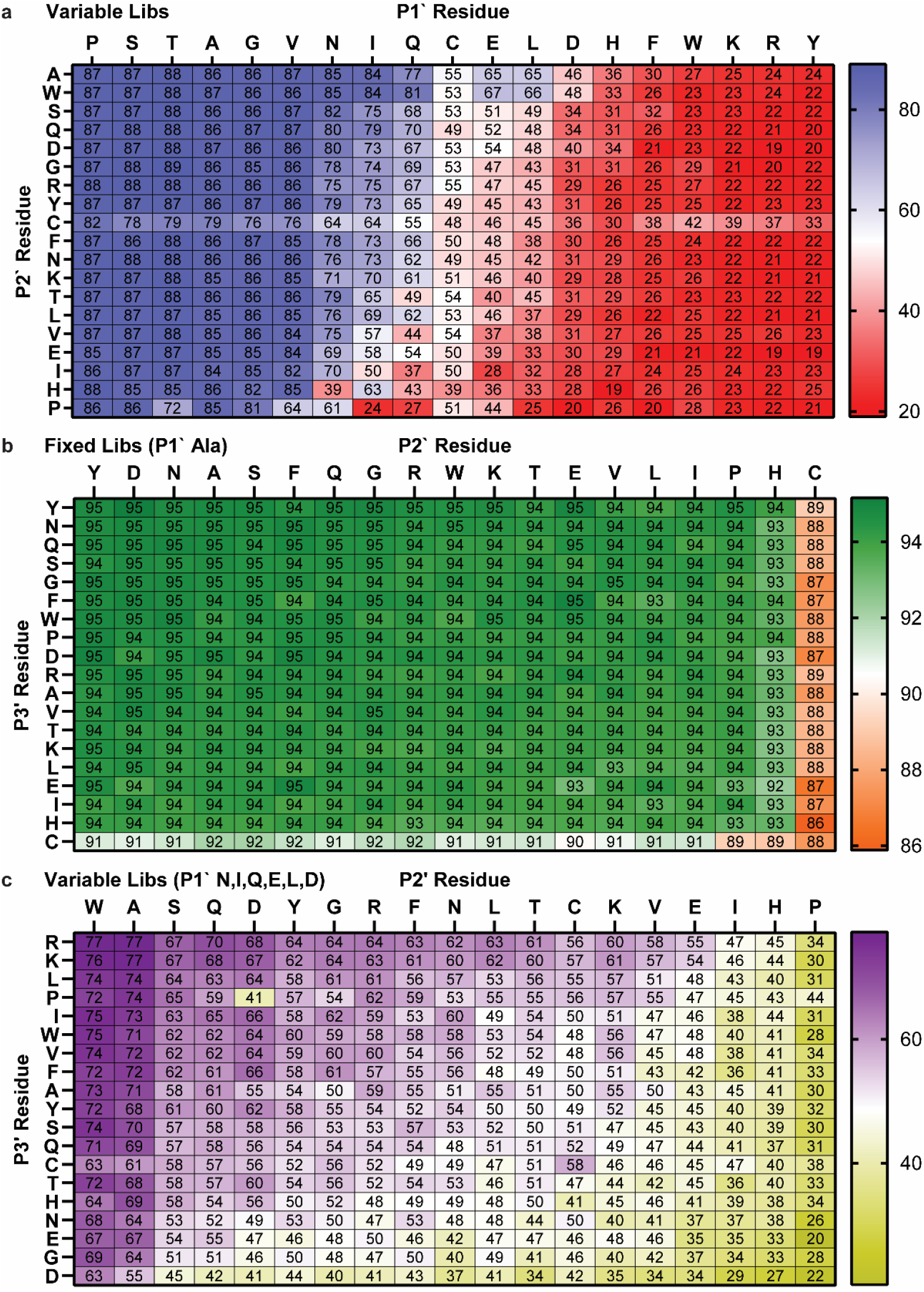
Heatmaps of percent cleavage calculated from subgroup analyses of two-residue combinations: P1’ × P2’ using the averaged Variable libraries (a), P2’ × P3’ using averaged Fixed libraries (fixed P1’ Ala) (b), and P2’ × P3’ using averaged Variable libraries (fixed P1’ Asn, Ile, Gln, Glu, Leu, or Asp) (c). Columns and rows are sorted by average values.

Subgroup analyses with combination P2’ × P3’ with constant P1’ position yielded some additional observations. The first analysis with the Fixed libraries confirms the known phenomenon that cleavage works well with Ala in the P1’ for most sequences, although the data suggest again that Cys interferes with cleavage slightly at either the P2’ or P3’ position (**Fig. 6b, correlation in Supplementary Fig. S11b**). Similar subgroup analyses using the portions of variable libraries filtered and analyzed separately for fixed Ser, Thr, and Ala in the P1’ position yielded similar results (**Supplementary Fig. S12**). For these P1’ Tier 1 amino acids, sequences cleaved efficiently with any amino acid in the P2’ and P3’ positions except for Pro, Cys, His, and a few other amino acids that varied according to the specific amino acid in the P1’ position. To best discern the impact of amino acid residues at the P2’ and P3’ positions, an analogous heatmap was generated from sequences with a Tier 2 P1’ amino acid (Asn, Ile, Gln, Asp, Leu, or Asp) (**Fig. 6c, correlation in Supplementary Fig. S11c**). This analysis suggests that Trp and Ala are advantageous in the P2’ position, allowing for relatively efficient cleavage for all amino acids in the P3’ position, with basic amino acids Lys and Arg slightly preferred. Pro, His, Ile, and Glu were generally disfavored in the P2’ position and Asp, Glu, and Gly were disfavored in the P3’ position.

### Comparison to Literature

In general, conclusions from selection data agree with previously published literature studying the cleavage efficiency of Met and HPG by MAP (summarized in **Supplementary Table S4**). The importance of side chain size of the amino acid in the P1’ position is universally acknowledged with previous literature concluding that smaller side chains favor cleavage and larger side chains disfavor cleavage. The exact same six P1’ Tier 1 amino acids were identified by Hirel *et al*.^9^, and all other previous literature list some subset of these 6 amino acids.^6–8,10–13^ Considering the more subtle impact of amino acid residues in downstream positions, there is generally less agreement concerning the P2’ position, with only Frottin *et al*.^6^ reporting that Trp favors cleavage and that both Pro and Glu disfavor cleavage. Frottin *et al*. also suggests that Cys in positions further downstream may disfavor cleavage efficiency. While some publications have reported a negative impact of acidic residues at the P3’ position, we are the first to detect a positive impact of basic residues Lys and Arg at that position.

### Kinetic Assay

To validate the selection results, a collection of heptapeptides with select P1’ and P2’ amino acid combinations was prepared by Fmoc solid-phase synthesis (**Supplementary Scheme S1**) and assayed for cleavage efficiency by an *in vitro* time-course kinetic experiment adapted from Merkel *et al*.^12^ All sequences were tested with both natural methionine (M) and HPG (X) at the N-terminus. An assay mixture containing enzyme and a large excess of substrate was incubated at 37 °C, and a small aliquot was removed at each time point and quenched with EDTA. Samples were then diluted with water containing HOBt as quantification standard and run on LC-MS with single ion recording (SIR). Percent cleavage was calculated from standardized area under the curve relative to the initial measurement. All assays were run in triplicate.

To visualize the data, percent cleavage values were plotted in Prism v10.4.2 (**Fig. 7a,b time-course in Supplementary Fig. S17**). Peptides starting with natural Met could be split into 4 tiers: 1) MAT, MAW, and MSW, which were approximately fully cleaved by 10 min, 2) MAE, MGT, and MSE, which were approximately fully cleaved by 20 min, 3) MAP, which was approximately fully cleaved by 130 min, and 4) MGP and MEP, which did not demonstrate any observable cleavage within the timeframe of the assay. Similarly, peptides with N-terminal HPG could be split into 3 tiers: 1) XAW and XSW, which were mostly cleaved by 36 min, 2) XAT, XAE, and XSE, which were mostly cleaved by 130 min, and 3) XGP and XAP, which did not demonstrate observable cleavage within the timeframe of the assay. In all cases, peptides with N-terminal HPG demonstrated lower cleavage efficiency than their analogous counterparts with N-terminal Met.

**Figure 7:**
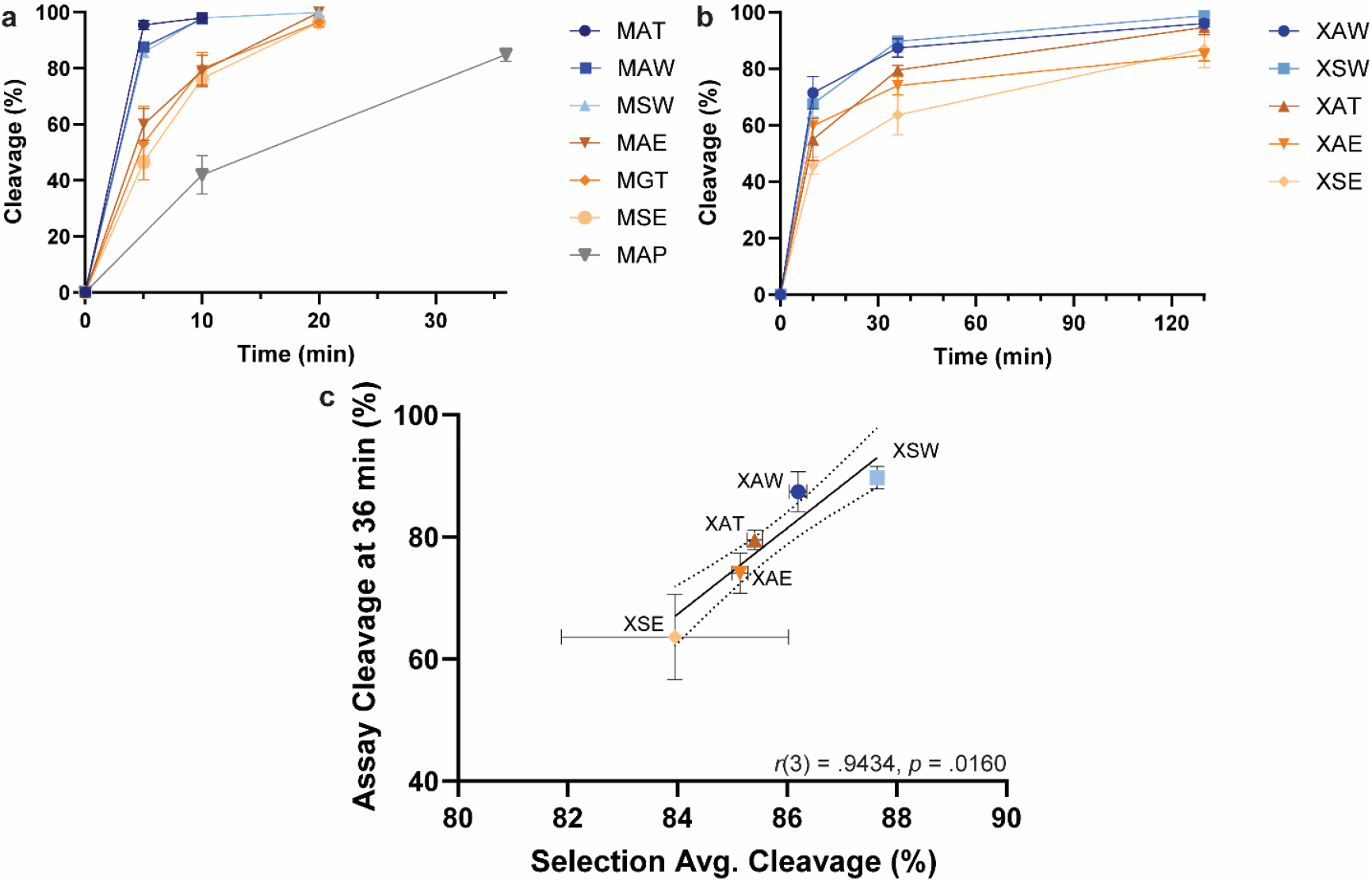
(a) Percent cleavage observed for synthetic heptapeptides treated with MAP and quantified by LC/MS (triplicate data). Peptides start with the sequence displayed (including N-terminal Met), followed by TYNK (b) Experiment analogous to (a) with N-terminal HPG. (c) Correlation of synthetic peptide cleavage assay with percent cleavage of peptides in selection. Selection data are averaged for peptides beginning with sequence shown, followed by Thr in the P3’ position and all random amino acids subsequently. Vertical and horizontal error bars represent standard deviation, and the Pearson correlation coefficient is given. A linear regression best-fit line is given with dotted lines representing 95% confidence bands. Colors correspond to cleavage efficiency tiers as in **Figure 4** (blue = Tier 1; orange/brown = Tier 2; gray = Tier 3).

For comparison of results from kinetic assay and selection data, percent cleavage at a single time point from the kinetic assay (36 min) was plotted against the appropriate subgroups from analysis with combination P1’ × P2’ with fixed Thr in the P3’ position (full heatmap in **Supplementary Fig. S18**). The results demonstrated moderate correlation (**Fig. 7c**). It is important to note the caveat that all synthetic peptides tested in the kinetic study contained a single sequence TYNK at positions P3’-P6’, which was designed for UV and MS detection sensitivity, but may not have been representative of average trends observed in the selection.

Taken together, our kinetic study suggests the following cleavage efficiency trends at each given position: 1) P1: Met > HPG; P1’: Ala ≅ Ser > Gly; P2’: Trp > *Thr > Glu >> Pro, with the caveat that MAT (but not XAT) cleaved more efficiently than analogous peptides with Trp in the P2’ position.

## Conclusion

In summary, we have utilized mRNA display and NGS to profile the substrate specificity of MAP with unnatural peptides containing N-terminal HPG. Analysis of cleavage data across various subgroups of peptides indicated a strong preference for cleavage with small and nonacidic residues in the P1’ position, a preference for Trp and Ala and aversion for Pro, His, and Ile in the P2’ position, a weak preference for basic residues and aversion for acidic residues in the P3’ position, potentially a weak aversion to acidic residues in the P4’ position, and no discernable trends at positions further downstream except for an aversion for Cys. Several of these observations could be rationalized by predicted enzyme-substrate complex modeling. To validate selection results, we chemically synthesized several heptapeptides and conducted time-course kinetic assays whose results correlated with selection data. It is important to note that while we designed the selection experiment to correct for variations in CuAAC biotinylation and pulldown efficiency, it is possible that over/undercorrection may affect the results, especially for cysteine-containing peptides where the correction was large. While our results correspond well with previous literature using both N-terminal Met and HPG, the assay could potentially be redesigned with repeated rounds of selection to increase signal/noise, or with natural, biologically relevant Met using novel redox-based Met-specific bioconjugation.^34^ These results may assist in the design of peptides and proteins with better control over cleavage of N-terminal Met or HPG.

## Supporting Information

Additional experimental details, figures, tables, and data including materials, methods, primer sequences, scanned SDS-PAGE gels, individual and select additional subgroup analyses, correlation and regression analyses, additional AlphaFold predicted models, peptide synthetic scheme, and representative LC-MS kinetic assay data.

## Acknowledgments

This work was supported by NIH grants R01-AI113737 and R01-AI090745. The graphical abstract was created with icons from BioRender. *Turra, R. (2025) https://BioRender.com/0ygjtvx*.

## TOC Graphic

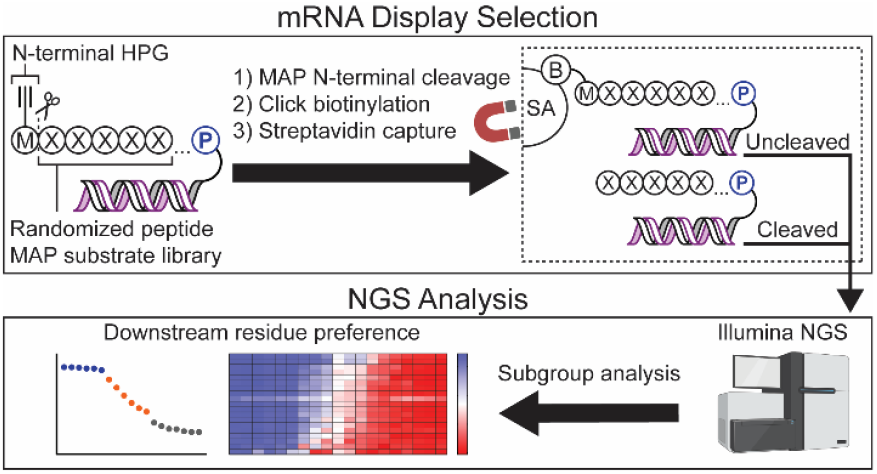

