## Supporting Information for "Systematic profiling of peptide substrate specificity in N-terminal processing by methionine aminopeptidase using mRNA display and an unnatural methionine analogue"

### I. mRNA Display Selection

#### A. Materials

Synthetic short oligos and ssDNA libraries (**Table S1**) were purchased from W.M. Keck Oligonucleotide Synthesis Facility (Keck) or Integrated DNA Technologies (IDT) and if over 50 nt in length gel purified by denaturing urea-PAGE gel (SequaGel, National Diagnostics; CBS vertical electrophoresis system, 16.5 cm × 14.5 cm). XL-PSO was gel purified by NAP25 column and eluted into water. dNTP and NTP sets were purchased from Thermo Scientific. Nuclease P1 was purchased from New England Biolabs (NEB). Reagents and buffers, usually molecular biology grade, were purchased from Thermo Scientific or Sigma, and homemade buffers were generally prepared in MilliQ Ultrapure or RNA grade water (Fisherbrand) and sterilized by 0.22 µm PES filters (Millipore Stericup). Biotin-$PEG_{11}$-azide was purchased from Alfa Aesar. $Man_9$-cyclohexyl-azide and tris(3-hydroxypropyltriazolylmethyl)-amine (THPTA) were prepared in-house.[1] C1000 Touch Thermal Cycler (Biorad), NanoDrop One$^{C}$ (Thermo Scientific), and Model 422 Electro-Eluter (Biorad) were used for PCR, quantitation, and electroelution, respectively. SDS-PAGE gels were either homemade from Bis-Acrylamide 37.5:1 (Fisher Bioreagents) or purchased precast (4-20% gradient, Biorad). L-Histidine, [ring-2,5-$^{3}$H]- (Moravek Biochemicals) was used to radiolabel translated peptides. Tritium-containing peptides were visualized by fluorography using Amersham Amplify Fluorographic Reagent (Cytiva), Blue Devil Premium autoradiography film (Genesee Scientific), and Konica Minolta SRX-101A processor. Liquid scintillation counting was performed on a Beckman Coulter LS6000TA scintillation counter using Econo-Safe Biodegradable Counting Cocktail (Atlantic Nuclear) in 20 mL HDPE scintillation vials with PP cap (Fisherbrand). MALDI spectra were acquired on a Voyager DE-PRO MALDI-TOF instrument using α-cyano-4-hydroxycinnamic acid matrix (CHCA; Sigma). PURE system enzymes, MAP, and PDF were expressed in-house.[2,3]

**Table S1**: Library and primer sequences used in selection. All were ordered from IDT except for Long Libraries and XL-PSO which were ordered from Keck. HPG is denoted by " *M* ".

| Name | Sequence 5` to 3` |
|---|---|
| Variable-Long | TAATACGACTCACTATAGGGTTAACTTTAGTAAGGAGGACAGCTAAATGNNSNNSNNSNNS<br>NNSNNSNNSNNSNNSGGCTCCGGTAGCTTAGGCCACCATCACCATCACCACCGGCTATAGG<br>TAGCTAG |
| | *M*XXXXXXXXXGSGSLGHHHHHHRL |
| Fixed-Long | TAATACGACTCACTATAGGGTTAACTTTAGTAAGGAGGACAGCTAAATGGCGNNSNNSNNS<br>NNSNNSNNSNNSNNSGGCTCCGGTTCTCTGGGTCATCACCACCATCACCACCGGCTATAGG<br>TAGCTAG |
| | *M*AXXXXXXXXGSGSLGHHHHHHRL |
| Variable-Short | TAATACGACTCACTATAGGGTTAACTTTAGTAAGGAGGACAGCTAAATGNNSNNSNNSNNS<br>NNSGGCTCCGGTAGCTTAGGCCACCATCACCATCACCACCGGCTATAGGTAGCTAG |
| | *M*XXXXXGSGSLGHHHHHHRL |
| Fixed-Short | TAATACGACTCACTATAGGGTTAACTTTAGTAAGGAGGACAGCTAAATGGCGNNSNNSNNS<br>NNSGGCTCCGGTTCTCTGGGTCATCACCACCATCACCACCGGCTATAGGTAGCTAG |
| | *M*AXXXXGSGSLGHHHHHHRL |
| T7 Primer | TAATACGACTCACTATAGGGTTAACTTTAG |
| XL-PSO | (C6 psoralen)-**UAGCCGGUG**(dA)$_{15}$(Spacer 9)$_2$dA(dC)$_2$-Puromycin<br>(**Bold = 2′-OMe nucleotides**) |
| PCB18-RT-primer1 | /5PCBio//iSp18/TTTTTTTTTTTTTTTGTGATGGTGATGGTGGCCTAAGC |
| PCB18-RT-primer2 | /5PCBio//iSp18/TTTTTTTTTTTTTTTGTGATGGTGGTGATGACCCAGAG |
| MAP-Lib-FP | TAATACGACTCACTATAGGGTTAACTTTAGTAAGGAGG |
| MAP-Var-RP | CTAGCTACCTATAGCCGGTGGTGATGGTGATGGTGGCCTAAGC |
| MAP-Fix-RP | CTAGCTACCTATAGCCGGTGGTGATGGTGGTGATGACCCAGAG |

**B. Selection**

*1. Preparation of Library DNA*
The antisense strands of DNA libraries were purchased either from Keck (Long libraries, 200 μmol scale) or from IDT (Short libraries, Ultramer® DNA Oligos, 4 nmol scale). Lyophilized libraries were dissolved in 250 μL water, and to a 50 μL portion of resuspended libraries was added 50 μL 8 M urea. Libraries were purified by 8% urea-PAGE gel, eluted into 1 mL 0.4 M KOAc (pH 5.5), isopropanol precipitated with 70% (v/v) ethanol rinse, and resuspended in 21 μL water. Yields were between 800 and 1200 pmol for Long libraries and between 260 and 300 pmol for Short libraries.

*2. Preparation of Library RNA*
Purified ssDNA libraries (50 pmol) were transcribed by T7 transcription reaction (250 μL) with 1.2 equivalent of a DNA primer containing the T7 promoter sequence. T7 transcription reactions contained final concentrations of 80 mM HEPES-KOH (pH 7.6), 40 mM DTT, 2 mM spermidine, 25 mM $MgCl_2$, 50 mg/mL PEG-8000, 4 mM GTP, 4 mM UTP, 4 mM ATP, 4 mM CTP, 2.5 U/mL inorganic pyrophosphatase, 240 nM T7 primer, 0.05 mg/mL homemade T7 RNA polymerase, and 200 nM ssDNA library. After overnight incubation at 37 °C, crude transcripts were incubated with a final concentration of 0.05 U/μL Turbo DNase (Invitrogen) at 37 °C for 15 minutes to remove template DNA before quenching the DNase with EDTA (pH 8.0) at a final concentration of 37.5 mM. RNA libraries were ethanol precipitated, resuspended in 100 μL 8 M urea, and purified by 8% denaturing urea-PAGE on plates presoaked in NaOH to prevent contamination. Excised gel slices were electroeluted using Bio-Rad Model 422 Electro-Eluter. Recovered samples (700 – 850 μL) were precipitated by isopropanol (using 6% volume 5 M NaCl salt) followed by 70% (v/v) ethanol rinse. RNA library pellets were resuspended in 100 μL water and quantitated by NanoDrop. Yields were between 2300 and 6900 pmol.

Purified RNA libraries (1215 pmol) were photo-crosslinked with 1.5 eq. of puromycin-containing oligo XL-PSO in crosslinking reactions (405 μL) containing final concentrations of 20 mM HEPES-KOH (pH 7.6), 0.1 M KCl, 1 mM spermidine, 1 mM EDTA, 7.5 μM XL-PSO, and 3 μM RNA library. Reaction mixtures were split into 8-strip 0.2 mL PCR tubes in 50 μL aliquots for annealing on a thermal cycler by an incubation cycle of 70 °C for 3 minutes, slow cooling to 25 °C (0.1 °C/sec), 25 °C for 5 minutes, and chill to 4 °C. After annealing, reaction mixtures were transferred to a Costar 96-well plate (100 μL/well) and irradiated for 20 min on ice at 365 nm using a handheld UV lamp. The puromycin-modified library RNAs were purified by precipitation with isopropanol followed by 70% ethanol rinse, 8% denaturing urea-PAGE with visualization by ethidium bromide staining (0.5 μg/mL), and finally electroelution. Successful isolation of crosslinked RNA was verified by running samples taken from the reaction mixture before UV irradiation, the crude product before gel purification, and the purified crosslinked RNA. Yields of crosslinked RNA libraries were between 175 and 250 pmol.

*3. Library Translation and mRNA-peptide Fusion Formation*
Purified crosslinked RNA libraries (175 pmol) were subjected to PURE system translation reactions to form mRNA-peptide fusions. Translation reactions (350 μL) contained final concentrations of 50 mM HEPES-KOH, pH 7.6, 12 mM magnesium acetate, 2 mM spermidine, 100 mM potassium glutamate, 1 mM dithiothreitol (DTT), 1X cOmplete ULTRA, EDTA-free (Roche), 1 mM ATP, 1 mM GTP, 20 mM creatine phosphate (Calbiochem), 48 $ABS_{260}$ tRNA from *E. coli* MRE 600 (Roche), 10 μg/mL 5,10-methyltetrahydrofolate, 0.04 $ABS_{280}$ creatine kinase (Roche), 0.85 U/mL nucleoside 5′-diphosphate kinase from bovine liver (Sigma), 6.8 U/mL myokinase from rabbit muscle (Sigma), 1 U/mL inorganic pyrophosphatase, 20 μg/mL MTF, 10 μg/mL IF1, 40 μg/mL IF2, 10 μg/mL IF3, 100 μg/mL EF-Tu, 50 μg/mL EF-Ts, 50 μg/mL EF-G, 10 μg/mL RF1, 10 μg/mL RF3, 10 μg/mL RRF, 0.66 μM MetRS, 0.23 μM GluRS, 0.027 μM PheRS, 0.21 μM AspRS, 0.45 μM SerRS, 0.011 μM ThrRS, 0.021 μM ArgRS, 0.27 μM GlnRS, 0.11 μM IleRS, 0.093 μM LeuRS, 0.23 μM TrpRS, 0.094 μM AsnRS, 0.21 μM HisRS, 0.18 μM TyrRS, 0.089 μM ValRS, 0.031 μM ProRS, 0.070 μM AlaRS, 0.41 μM CysRS, 0.18 μM LysRS, 0.024 μM GlyRS, 1.2 μM ribosomes, a mixture of 17 natural amino acids (0.3 mM each), with methionine, cysteine, and histidine omitted and preadjusted pH to 7.6 with KOH. To these translation reactions were added to final concentrations of 0.3 mM cysteine, 0.3 mM L-homopropargylglycine (Chiralix), 3.1 μM [2,5-$^{3}$H]-L-histidine (Moravek Biochemicals) to radiolabel peptides, and 0.5 μM crosslinked RNA. Translation reactions were incubated at 37 °C for 30 minutes before adding 0.3 volume of a Mg/K solution containing 172 mM magnesium acetate and 2.05 M potassium chloride and letting sit at room temperature for an additional 15 minutes. Crude translation samples were then stored overnight at -20 °C to stabilize mRNA-peptide fusion formation.

*4. Library Purification, Reverse Transcription, and PDF Digestion on Streptavidin Resin*

mRNA-peptide fusion libraries were purified and reverse transcribed using Pierce™ Streptavidin UltraLink™ Resin (SAUR; Thermo Scientific) as represented in **Fig. S1** as follows: Unless otherwise stated, all washes were performed with 500 µL buffer. For 300 µL translation mixture of each library, 60 µL (0.2 vol.) of the resin was washed 3 times with 300 µL SAUR Washing Buffer 1 (SWB1 Buffer; 50 mM Tris-HCl, pH 8.0, 150 mM KCl, 0.2% (v/v) Triton X-100, 7 mM BME) and then resuspended in 375 µL of a fusion/primer mix comprising 300 µL translation mixture, 2.5 mM Tris-HCl, pH 8.0, 60 mM EDTA, pH 8.0, 0.2% (v/v) Triton X-100, and 0.56 µM PCB (photocleavable biotin)-18-RT-primer (PCB18-RT-primer2 for Fixed libraries, PCB18-RT-primer1 for the Variable libraries). After tumbling for 30 minutes at room temperature, resins were spun down and washed twice with SWB1 Buffer (second wash 1 mL). To perform the reverse transcription, the washed resins were heated to 65 °C for 10 minutes (without adding buffer) and then chilled on ice for at least 10 minutes. Resins were resuspended in 90 µL (1.5 resin vol.) of a reverse transcription reaction mix (1X Reaction Buffer (Thermo Scientific), 1 mM of each dNTP, 1 U/µL RiboLock RNase Inhibitor (Thermo Scientific), 10 U/µL RevertAid H minus Reverse Transcriptase (Thermo Scientific)) followed by incubation at 42 °C for 1 hour with occasional mixing. For PDF digestion, resins were washed twice with PM buffer (50 mM Tris-HCl, pH 7.8, 0.1 mM $CoCl_2$, 0.1% (v/v) Triton X-100), resuspended in 90 µL PM buffer containing 6.5 µM PDF, and tumbled for 1 hour at 37 °C. The resins were washed 3 times with SWB1 using a volume of 1 mL for the third wash and resuspended in 120 µL (2 resin vol.) of SWB1. To photo-elute cDNA/mRNA-peptide fusions from the resin, the tubes were covered on the side with aluminum foil and irradiated from the top by a 365 nm handheld UV lamp for 20 min. The supernatant and an equal volume rinse were collected before repeating the irradiation for another 10 min followed by another supernatant collection and rinse.

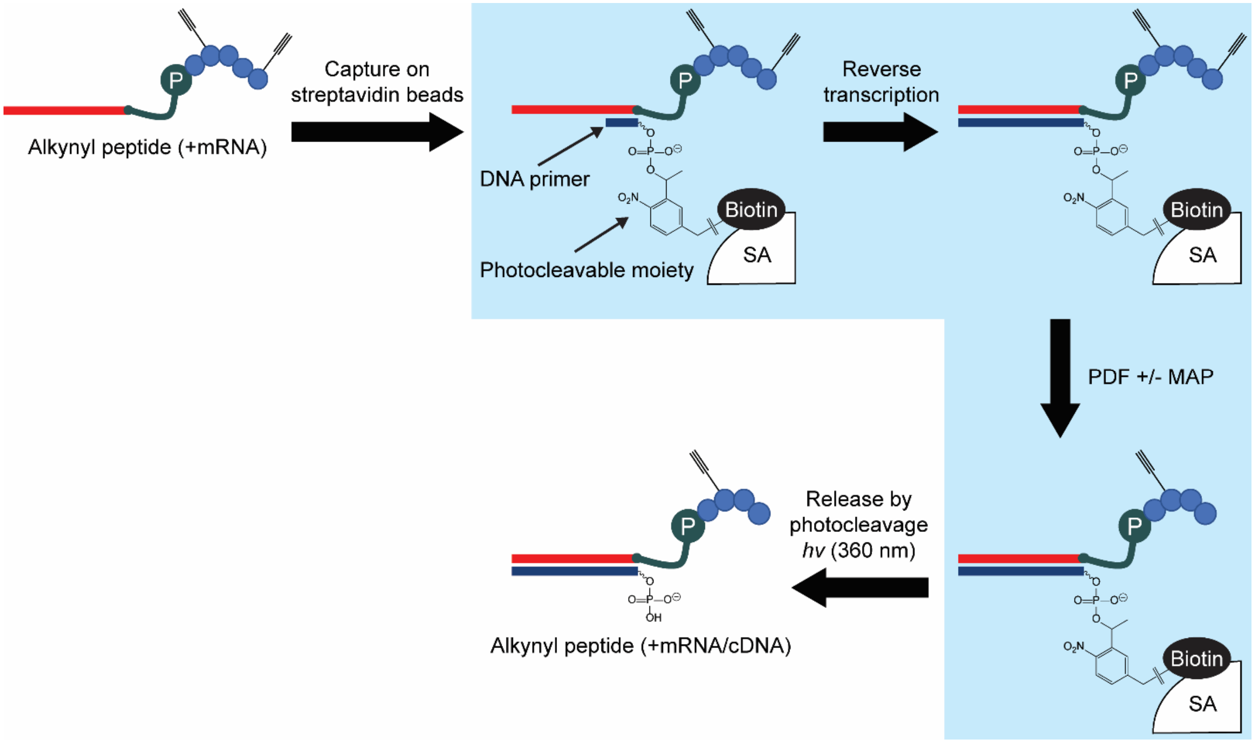

**Figure S1**: Workflow for capture on streptavidin Ultralink™ resin via photocleavable biotinylated reverse transcription primers, sequential enzymatic reactions performed on-resin, and recovery after photocleavage.

*5. Co-translational vs Post-translational MAP Digestion*
Although we initially planned to perform PDF and MAP digestions co-translationally, we discovered that the mRNA displayed short peptide libraries used in this selection were not compatible with co-translational MAP cleavage. During mRNA-peptide fusion formation, the ribosome stalls at the stop codon, allowing puromycin to be added to the end of the nascent peptide. We promote ribosome stability and puromycin incorporation by adding $Mg^{2+}$ and $K^{+}$ and incubating for 15 min at room temperature before keeping at -20 °C overnight and purification the next day with EDTA-containing buffers that encourage ribosome dissociation. Since the 20- or 24-mer nascent peptide is contained completely within the ribosomal exit tunnel, which proteolytic cleavage experiments and cryo-EM structures of *E. coli* ribosomes have confirmed covers approximately 30-40 amino acids,[4,5] the peptide N-terminus would be inaccessible for cleavage by PDF or MAP. Furthermore, since the EDTA used in purification would also chelate the necessary cobalt cofactor necessary for PDF and MAP activity, the enzymes would still be unable to digest the peptides even after ribosomal dissociation.

To test this hypothesis, we prepared four samples of the Fixed-Long library (with and without co-translational MAP digestion, each with and without post-translational MAP digestion) and subjected them to click reaction with $Man_9$-cyclohexyl-azide, a large glycan used in previous mRNA display projects, to assess whether the N-terminal HPG had been cleaved or not. After nuclease P1 digestion and SDS-PAGE, we concluded that post-translational digestion successfully removed the alkyne while co-translational digestion did not (**Fig. S2**).

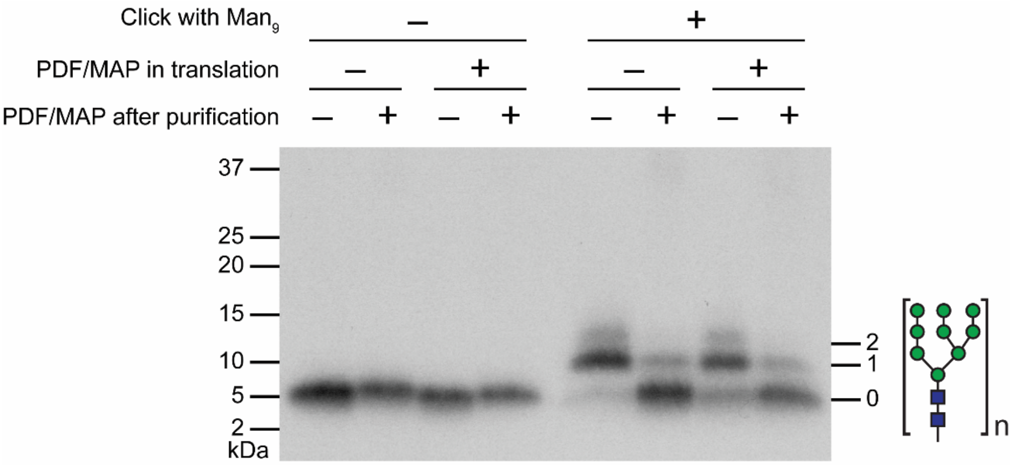

**Figure S2**: Comparison of PDF/MAP processing during and after library translation. Fixed-Long fusion library was prepared with or without PDF/MAP in translation, and both versions were purified with or without post-translational PDF/MAP digestion. Samples were clicked with $Man_9$-cyclohexyl-azide and results were visualized after nuclease P1 digestion by 4-20% SDS-PAGE and fluorography.

*6. Ni-NTA Agarose Purification*
To remove cDNA/mRNA duplexes not fused to peptide, cDNA/mRNA-peptide fusions were purified by immobilized metal affinity chromatography (IMAC) as follows: cDNA/mRNA-peptide fusions were buffer exchanged by precipitation with isopropanol in the presence of 10 µg/microcentrifuge tube linear acrylamide carrier (Invitrogen) followed by 70% (v/v) ethanol rinse and resuspension in 150 µL Denaturing Bind Buffer (DBB; 100 mM $NaH_2PO_4$, 10 mM Tris, 6 M guanidinium hydrochloride, NaOH to pH 8.0, 0.2% (v/v) Triton X-100, 5 mM BME). 50 µL of 50% slurry HisPur™ Ni-NTA Resin for each library were washed twice with 400 µL DBB and resuspended in 50 µL DBB. Resin was transferred to fusions with an additional 50 µL DBB wash of the resin tube. After tumbling at 4 °C for 3 hours, ~250 µL of suspensions were transferred to a 0.22 µm Ultrafree-MC Centrifugal Filter (MilliporeSigma). The tubes containing the resin suspensions were rinsed with an additional 100 µL DBB transferred to the centrifugal filter unit. After spinning down at 2,000 ×g for 30 seconds, the resins were washed twice with 100 µL DBB, washed 3 times with 100 µL Native Wash Buffer (100 mM $NaH_2PO_4$, 300 mM NaCl, NaOH to pH 8.0, 0.2% (v/v) Triton X-100, 5 mM BME), and eluted 3 times with 50 µL Native Elute Buffer (50 mM $NaH_2PO_4$, 300 mM NaCl, 250 mM imidazole, NaOH to pH 8.0, 0.2% (v/v) Triton X-100, 5 mM BME). The final wash and each elution (after sitting at room temperature for 2 minutes) were spun down at 10,000 ×g for 30 seconds. Elution yields were quantitated by LSC indicating yields between 2.9 and 6.0 pmol.

*7. Gel Filtration*
To remove imidazole from Ni-NTA purified libraries, all 3 elution fractions were loaded onto a NAP5 column (GE Healthcare/Cytiva) preequilibrated with 10 mL of gel filtration buffer (10 mM Tris-HCl, pH 7.5, 1 mM EDTA, pH 8.0, 0.2% (v/v) Triton X-100, and 5 mM BME). 350 µL of gel filtration buffer was added for a total 500 µL flowthrough, and libraries were eluted in 700 µL gel filtration buffer.

*8. Negative Selection*
A negative selection was performed against the Dynabeads M-280 Streptavidin (Invitrogen) that are used in positive selection to remove library members that bound to the immobilization carrier. Libraries were isopropanol precipitated in the presence of 10 µg/microcentrifuge tube linear acrylamide carrier (Invitrogen) followed by 70% (v/v) ethanol rinse and resuspension in 50 µL SMBBv1 (20 mM Tris-HCl, pH 7.5, 500 mM NaCl, 0.1% (v/v) Triton X-100). To libraries were added 50 µL streptavidin beads washed three times with 250 µL SMBBv1, and the mixture was tumbled for 1 hour at room temperature. The supernatant was recovered to continue selection while beads were washed three times with 200 µL SMBBv1, heated at 95 °C for 5 min in 100 µL denaturing elute buffer (10 mM Tris-HCl, pH 7.5, 1 mM EDTA pH 8.0, 0.2% (w/v) SDS), washed with another 100 µL denaturing elute buffer, and resuspended in a third portion of 100 µL denaturing elute buffer for scintillation counting. Scintillation counting of supernatant, washes, and elutions indicated fraction bound values between 0.49 and 0.89%.

*9. Click Reaction*
Fusion libraries were ethanol precipitated in the presence of 10 µg linear acrylamide carrier (Invitrogen) per microcentrifuge tube and resuspended in 6 µL Mixture A (200 mM HEPES-KOH, pH 7.6, 10 mM aminoguanidine, 0.06% (v/v) Triton X-100). Samples were placed inside two-neck flasks alongside tubes of Mixture B (6 µL 12 mM Biotin-$PEG_{11}$-azide, 2 mM $CuSO_4$, 2 mM Tris(3-hydroxypropyltriazolylmethyl)amine ligand (THPTA)), Mixture C (7.2 µL 5 mM Biotin-$PEG_{11}$-azide, 0.833 mM THPTA), and a tube containing sodium ascorbate powder, with argon slowly going in one neck and passing out the other through a septum and needle. During this hour, the sodium ascorbate was dissolved in degassed water to 100 mM by removing the septum and inserting a pipette under Ar efflux. At the end of the hour, a pipette was used in the same way to transfer Mixture B and 1.2 µL sodium ascorbate to Mixture A for an approximately 12 µL reaction. After 15 min of Ar flow through the septum and needle, the needle was removed and tubes were rested for another 75 min under positive pressure. At this point Mixture C and 0.6 µL sodium ascorbate were transferred to the reaction mixture under Ar efflux, followed by another 15 min with Ar flow and another 75 min under positive pressure. The reaction was quenched with 3 µL 10 mM EDTA and diluted to 50 µL with gel filtration buffer. Libraries were then gel filtered by NAP5 column as in **Section I.B.7** to flowthrough volume of 500 µL but with elution in 750 µL gel filtration buffer.

*10. Round 1 Selection*
Libraries were isopropanol precipitated in the presence of 10 µg/microcentrifuge tube linear acrylamide carrier (Invitrogen) followed by 70% (v/v) ethanol rinse and resuspension in 10 µL SMBBv1 (+5 mM DTT). Since biotin-$PEG_{11}$-azide was dissolved in DMSO and represented a large portion of the click reaction volume, we were concerned that fusions could oxidize to form intermolecular disulfide bonds. Therefore, we added an incubation step with 5 mM DTT at 70 °C for 5 minutes before selection. Samples were then chilled on ice for at least 10 min and then diluted with SMBBv1 (-DTT) to a final volume of 50 µL.

To 50 µL Dynabeads M-280 Streptavidin (Invitrogen) beads prewashed in SMBBv1 (-DTT) and resuspended in 50 µL SMBBv1 (-DTT) were added 20 µL library and 30 uL SMBBv1 (+1 mM DTT), and the mixture was tumbled for 20 min at room temperature. The supernatant was recovered while beads were washed three times with 200 µL SMBBv1 (+0.5 mM DTT) and resuspended in 125 µL PCR Mix A (1X Phusion HF buffer, 0.2 mM each dNTP, 0.1% (v/v) Triton X-100). To measure the fraction bound, 12.5 µL beads were added to 100 µL denaturing elute buffer and heated at 95 °C for 5 min. The beads were then washed with another 100 µL denaturing elute buffer and resuspended in a third portion of 100 µL denaturing elute buffer for scintillation counting. From scintillation counting of supernatant, washes, elution, and resuspended beads we calculated fraction bound values given in **Table S2**. Before the selection, a small portion of the Variable-Short library was ethanol precipitated and subjected to a small-scale selection with varying incubation time, streptavidin or neutravidin coated beads, and concentrations of DTT to verify that none of these factors affected binding of biotinylated molecules to the streptavidin beads. A small portion of library was also subjected to a repeated click biotinylation followed by small-scale selection, where it was observed that clicking twice did not substantially improve bead uptake.

**Table S2**: Fraction bound values from selection given in percent.

| Digestion | Library | Round 1 | Round 2 |
|---|---|---|---|
| +PDF/-MAP | Var-Long | 38.07 | 47.02 |
| | Fix-Long | 49.32 | 52.63 |
| | Var-Short | 51.28 | 52.14 |
| | Fix-Short | 53.20 | 51.94 |
| +PDF/+MAP | Var-Long | - | 24.57 |
| | Fix-Long | - | 12.86 |
| | Var-Short | - | 23.50 |
| | Fix-Short | - | 5.86 |

*11. Recovery of Bound cDNA from Round 1 Selection*

DNA was recovered from the Round 1 bound fraction by on-bead PCR. To 90 µL beads remaining after pilot experiments was added 180 µL PCR Mix A and 90 µL PCR Mix B (1X Phusion HF buffer, 0.2 mM each dNTP, 4 µM forward primer (MAP-Lib-FP), 4 µM reverse primer (MAP-Var-RP for Variable Libraries; MAP-Fix-RP for Fixed Libraries), 0.08 U/µL Phusion Hot Start II DNA Polymerase (Thermo Scientific), 0.1% (v/v) Triton X-100) such that final concentrations were 1X Phusion HF buffer, 0.2 mM each dNTP, 1 µM forward primer, 1 µM reverse primer, 0.02 U/µL DNA polymerase, and 0.1% (v/v) Triton X-100. 24 µL PCR reactions were prepared and briefly vortexed to resuspend the beads just prior to placing on the thermal cycler running the following program: 98 °C for 30 sec, 7-9 cycles (Variable-Long: 9, Fixed-Long: 8, Variable-Short: 8, Fixed-Short: 7) of 98 °C for 5 seconds, 59 °C for 10 seconds, and 72 °C for 10 seconds.

PCR-amplified library DNA was isolated by removing the supernatant from magnetically isolated beads and then washing the beads once with 5 uL 0.1% (v/v) Triton X-100. Library DNA solutions were filtered by 0.22 µm Ultrafree-MC Centrifugal Filter (MilliporeSigma) to remove residual beads, extracted with phenol/chloroform and then chloroform, and precipitated with isopropanol followed by rinsing of pellets with 70% (v/v) ethanol. Yields were estimated by comparison to samples with known concentration on 8% native PAGE and determined to be between 9.1 and 16.0 pmol. For the next round of selection, a portion of the recovered DNA (3.8-6.7 pmol) was further amplified. An 80-fold dilution of recovered DNA into a 5 cycle PCR reaction provided enough material for subsequent experiments (97 to 107 pmol).

*12. Round 2 Selection*

For Round 2 of selection, cDNA-mRNA-peptide fusion libraries were prepared in the same way as in Round 1 with a few differences. Translation volume was increased to 488.56 µL (632.56 µL after addition of Mg/K solution) so libraries could be split into two 300 µL portions for purification and enzymatic reactions on streptavidin Ultralink™ resin. While one portion of each library was processed identically as in round 1, the other portion was processed with 15 µM MAP enzyme added to the digestion reaction with PDF.

Negative selection with M280 streptavidin magnetic beads was repeated a second time because some MAP-digested libraries had high fraction bound values. Fixed-Long and Fixed-Short libraries in particular had unusually high binding with 5.0 and 11.6% fraction bound, respectively. Because of this, a second negative selection with streptavidin beads was performed, in which fraction bound was below 0.5% in all libraries.

To further test the efficiency of the click reaction and verify that MAP digestion has occurred, we again used $Man_9$-cyclohexyl-azide as a click partner to enable visualization by SDS-PAGE. A small portion of each library was subjected to click reaction as in **Section I.B.9**, but with scaled down volumes (~5 µL reaction once Mixtures A and B mixed) and $Man_9$-cyclohexyl-azide at 10 mM (Mixture B) and 5 mM (Mixture C) instead of biotin-$PEG_{11}$-azide in each mixture. The libraries demonstrated the expected band pattern with the majority of non-MAP-digested libraries shifting up after successful click glycosylation and the majority of MAP-digested libraries remaining at baseline (**Fig. S3**). Since a small portion of each library may contain HPG within the randomized region, faint bands are visible indicating multiple click glycosylation events. It is also expected that Fixed libraries, with an Ala in the P1` position, demonstrate more complete cleavage such that a lower amount of clicked library is seen in MAP digested Fixed library samples than equivalent Variable library samples.

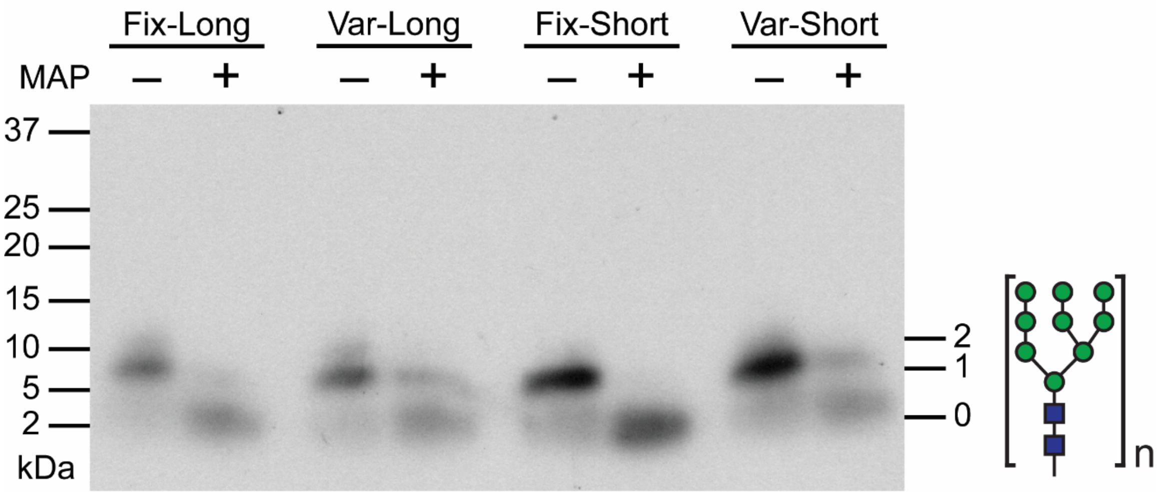

**Figure S3**: In preparation for Round 2 of selection, click reaction efficiency was tested across all +/- MAP treated libraries by click glycosylation with $Man_9$-cyclohexyl-azide to observe a gel shift in SDS-PAGE visualized by fluorography. Libraries demonstrated the expected band pattern with an upward shift in non-MAP-digested libraries, indicating successful conjugation of the glycan, and no upward shift in MAP-digested libraries, except for faint bands corresponding to poorer MAP cleavage of the Variable libraries.

Following the negative selection step, libraries were clicked and subjected to positive selection. The click reaction was conducted twice in the same manner as in Round 1 but with volumes scaled to reach ~10 µL reaction once Mixtures A and B were mixed. Libraries were prepared and partitioned with Dynabeads M-280 Streptavidin (Invitrogen) beads in the same manner as in Round 1 with all volumes scaled by 0.75X except for resuspension in PCR Mix A, which was increased to 200 µL. Fraction bound values are given in **Table S2**.

To probe whether all biotinylated library bound to the beads, a portion of library was subjected to selection, and the supernatant was used as the input for a second sequential selection step. Fraction bound values for this resubjection were very low (<1%), suggesting that under these selection conditions, the beads absorbed all of the biotinylated library.

*13. Preparation of DNA for NGS*

DNA was recovered from both bound and unbound fractions for each library. On-bead PCR was used to recover DNA from the bound fraction using the same protocol as in **Section I.B.11** but scaled volumes for 159 µL resuspended beads (for total PCR volume of 636 µL) and following a modified program: hold for 5 min at 98 °C, at which point samples were briefly vortexed to resuspend beads, 98 °C for 30 sec, 8-11 cycles (as determined by pilot PCR experiments) of 98 °C for 5 seconds, 59 °C for 10 seconds, and 72 °C for 1 min, followed by final extension at 72 °C for 5 min. PCR-amplified library DNA was collected and filtered in the same way as in **Section I.B.11**, with 5.0 to 20.0 pmol recovered. DNA was diluted 80-fold into a new PCR reaction (total 480 µL, 4 or 5 cycles) to further amplify DNA.

To recover DNA from the unbound fraction, the supernatant and three washes were gel filtered through a NAP5 column and eluted into 1000 µL gel filtration buffer. 947.5 µL of this elution was diluted by 1.4X into PCR reaction mixture (to the same final reagent concentrations as on-bead PCR, total PCR volume of 1326.5 µL) and run on the same PCR program as for on-bead PCR for 7-9 cycles (as determined by pilot PCR experiments).

Both bound and unbound fraction were then prepared for NGS. Recovered DNA libraries were purified by Monarch PCR and DNA Cleanup Kit (5 ug) (NEB). Samples were eluted in water and then adjusted to 10 mM Tris-HCl, pH 8.0. After quantitation by NanoDrop, libraries were combined into 4 pools (non-MAP-digested/bound, MAP-digested/bound, non-MAP-digested/unbound, MAP-digested/unbound) such that each pool contained the respective fraction of each of 4 starting libraries (Variable-Long, Fixed-Long, Variable-Short, Fixed-Short) at an equimolar ratio. Samples were submitted to GENEWIZ for sequencing on Illumina HiSeq (2 x 150 bp) with 50% PhiX spike-in for an estimated 7.3 million reads per library fraction.

### II. NGS Analysis

**Table S3**: Total number of reads and quantitation by scintillation counting for all 16 libraries sequenced by NGS and used in subgroup analyses.

| Digestion | Library | Fraction | Total reads | Quantitation (fmol) |
|---|---|---|---|---|
| +PDF/-MAP | Variable-Long | Bound | 6,009,695 | 188.6 |
| | | Unbound | 6,414,349 | 212.4 |
| | Fixed-Long | Bound | 5,681,420 | 181.1 |
| | | Unbound | 5,657,029 | 163.0 |
| | Variable-Short | Bound | 4,636,966 | 348.1 |
| | | Unbound | 5,036,631 | 319.5 |
| | Fixed-Short | Bound | 4,324,890 | 338.4 |
| | | Unbound | 4,666,361 | 313.0 |
| +PDF/+MAP | Variable-Long | Bound | 5,595,580 | 96.8 |
| | | Unbound | 7,223,715 | 297.2 |
| | Fixed-Long | Bound | 5,366,528 | 39.7 |
| | | Unbound | 6,547,605 | 269.3 |
| | Variable-Short | Bound | 4,346,871 | 115.0 |
| | | Unbound | 5,337,012 | 374.4 |
| | Fixed-Short | Bound | 3,932,730 | 26.0 |
| | | Unbound | 5,203,569 | 418.2 |

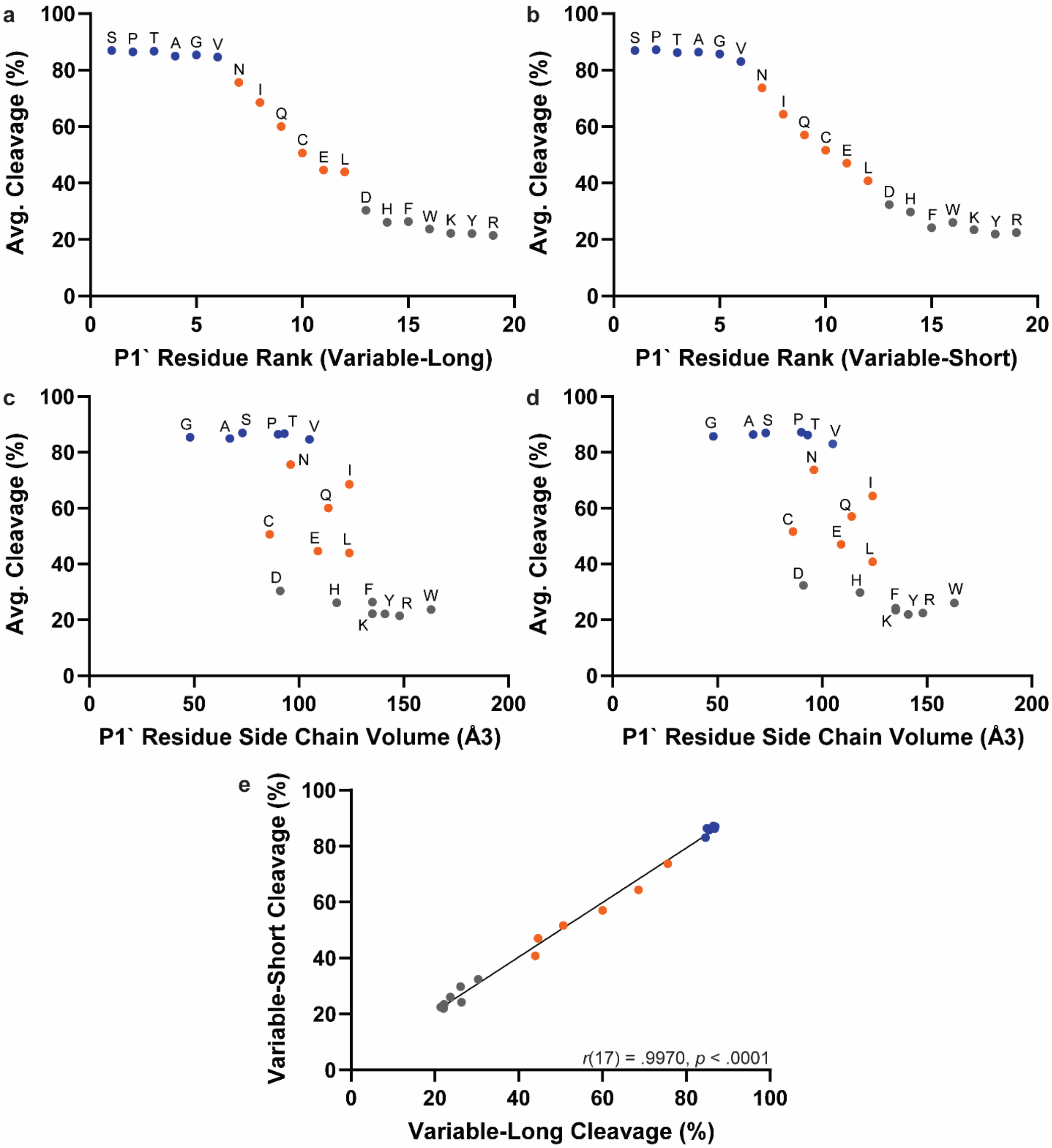

**Figure S4**: Estimated percent cleavage values from P1` subgroup analyses. (a-d) Percent cleavage values are presented for Variable-Long (a,c) and Variable-Short (b,d) libraries individually as a ranked list (a,b) and plotted against residue side chain size (c,d). (e) Variable-Long and Variable-Short library percent cleavage values are plotted against each other. The Pearson correlation coefficient is given, and the linear regression best-fit line is drawn to help visualize the linear relationship. Colors correspond to cleavage efficiency tiers as in **Figure 4** (blue = Tier 1; orange = Tier 2; gray = Tier 3).

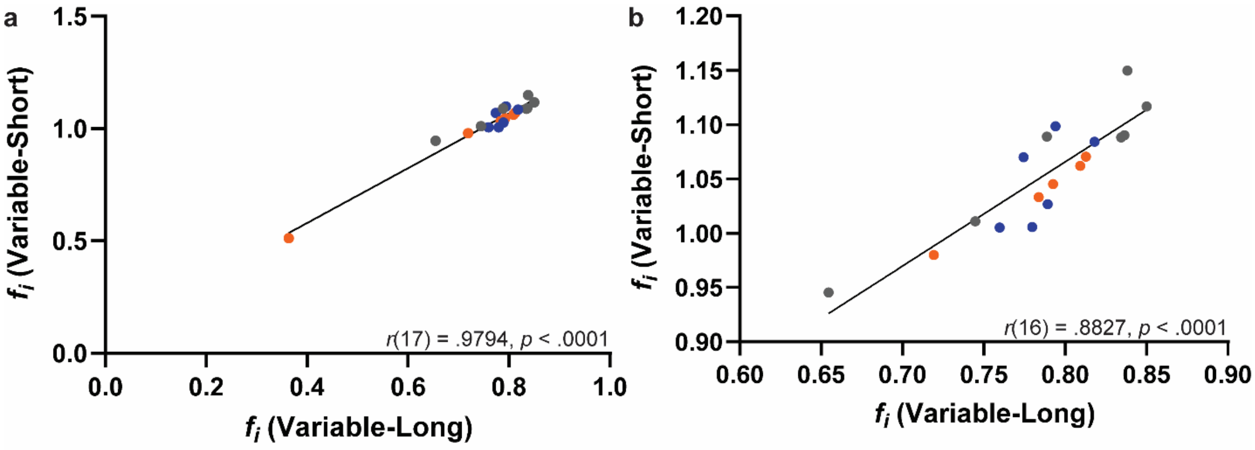

**Figure S5**: Click efficiency factor determined from P1` subgroup analysis for Variable-Long and Variable-Short libraries plotted against each other either including (a) or excluding (b) Cys. The Pearson correlation coefficient is given, and the linear regression best-fit line is drawn to help visualize the linear relationship. Colors correspond to cleavage efficiency tiers as in **Figure 4** (blue = Tier 1; orange = Tier 2; gray = Tier 3).

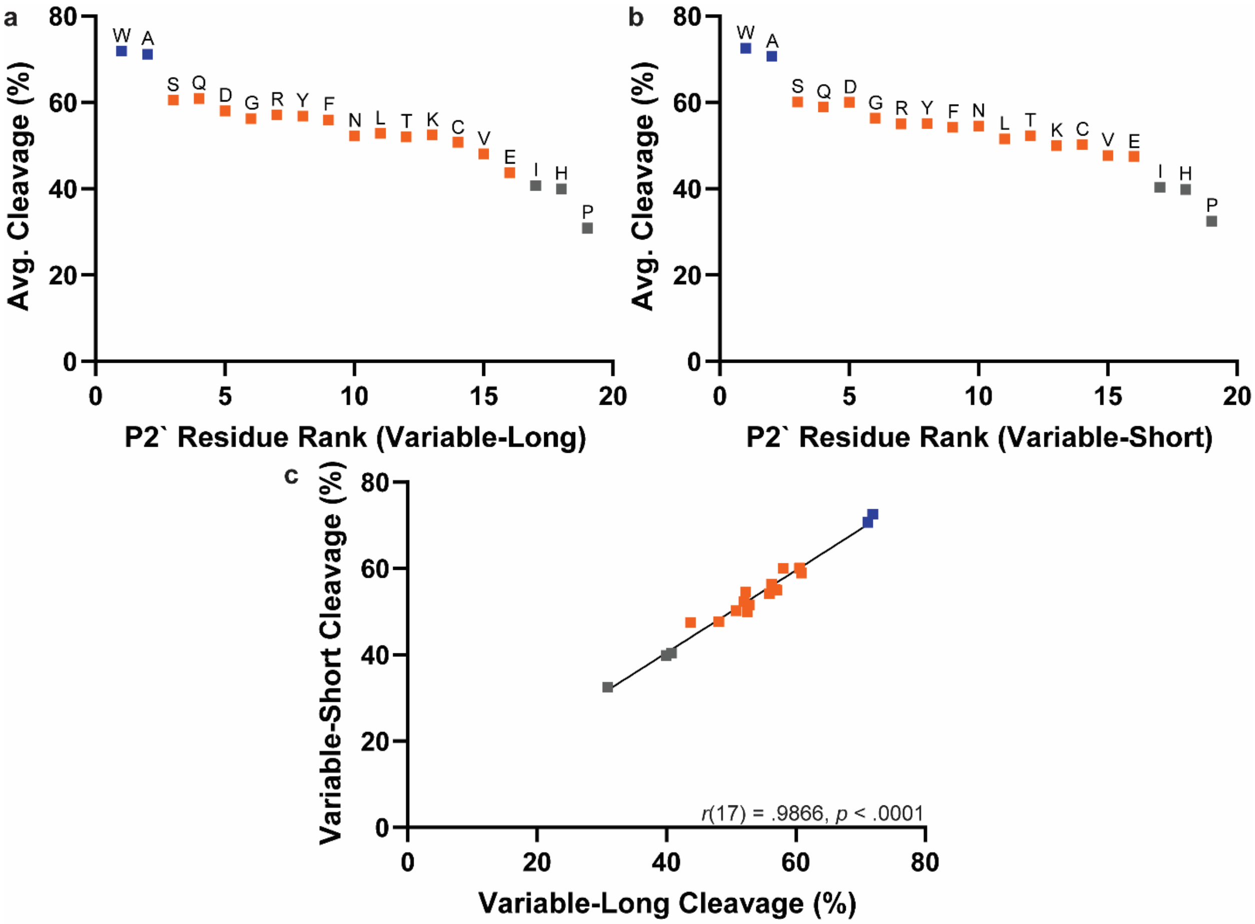

**Figure S6**: Estimated percent cleavage values from P2` subgroup analyses. (a-b) Percent cleavage values are presented for Variable-Long (a) and Variable-Short (b) libraries individually as a ranked list. (c) Variable-Long and Variable-Short library percent cleavage values are plotted against each other. The Pearson correlation coefficient is given, and the linear regression best-fit line is drawn to help visualize the linear relationship. Colors correspond to cleavage efficiency tiers as in **Figure 4** (blue = Tier 1; orange = Tier 2; gray = Tier 3).

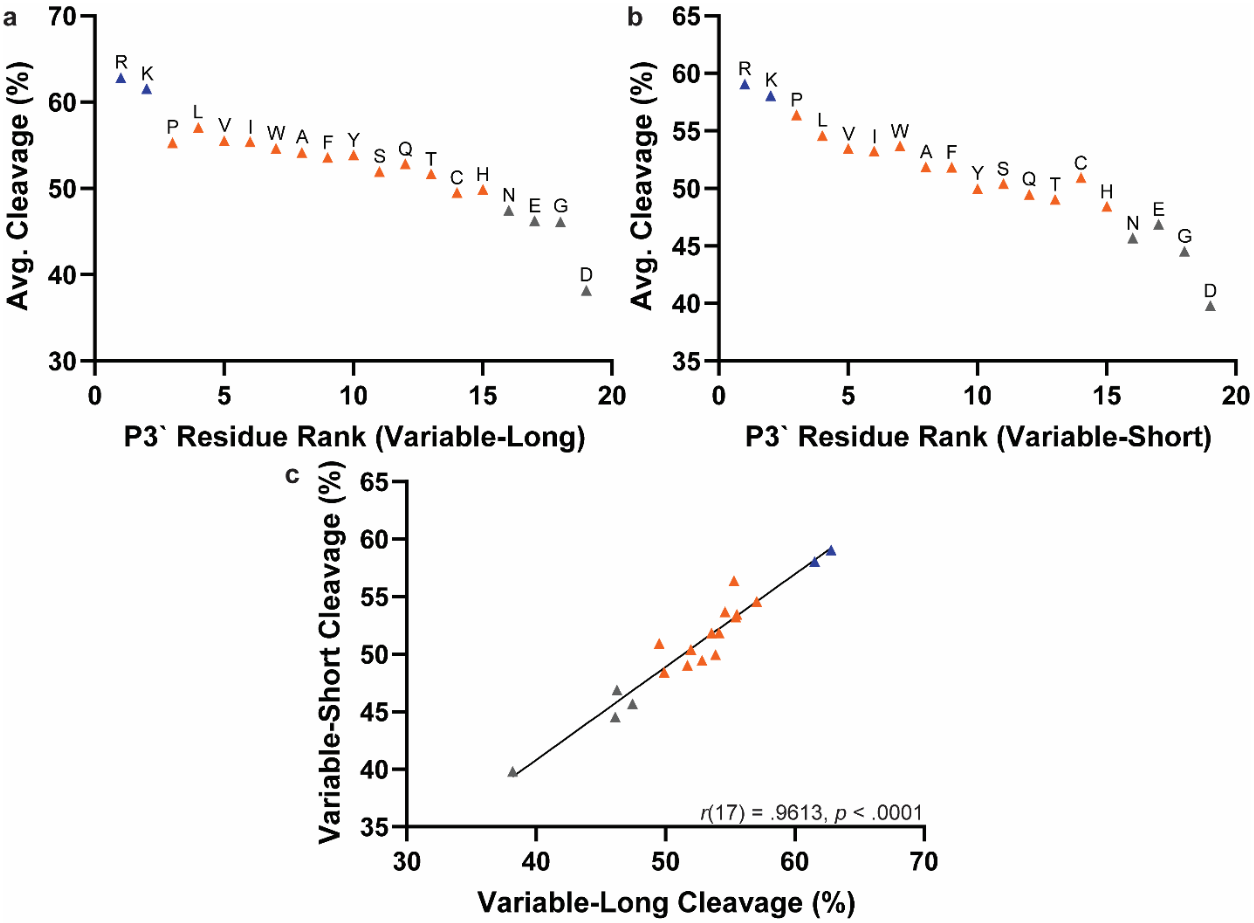

**Figure S7**: Estimated percent cleavage values from P3` subgroup analyses. (a-b) Percent cleavage values are presented for Variable-Long (a) and Variable-Short (b) libraries individually as a ranked list. (c) Variable-Long and Variable-Short library percent cleavage values are plotted against each other. The Pearson correlation coefficient is given, and the linear regression best-fit line is drawn to help visualize the linear relationship. Colors correspond to cleavage efficiency tiers as in **Figure 4** (blue = Tier 1; orange = Tier 2; gray = Tier 3).

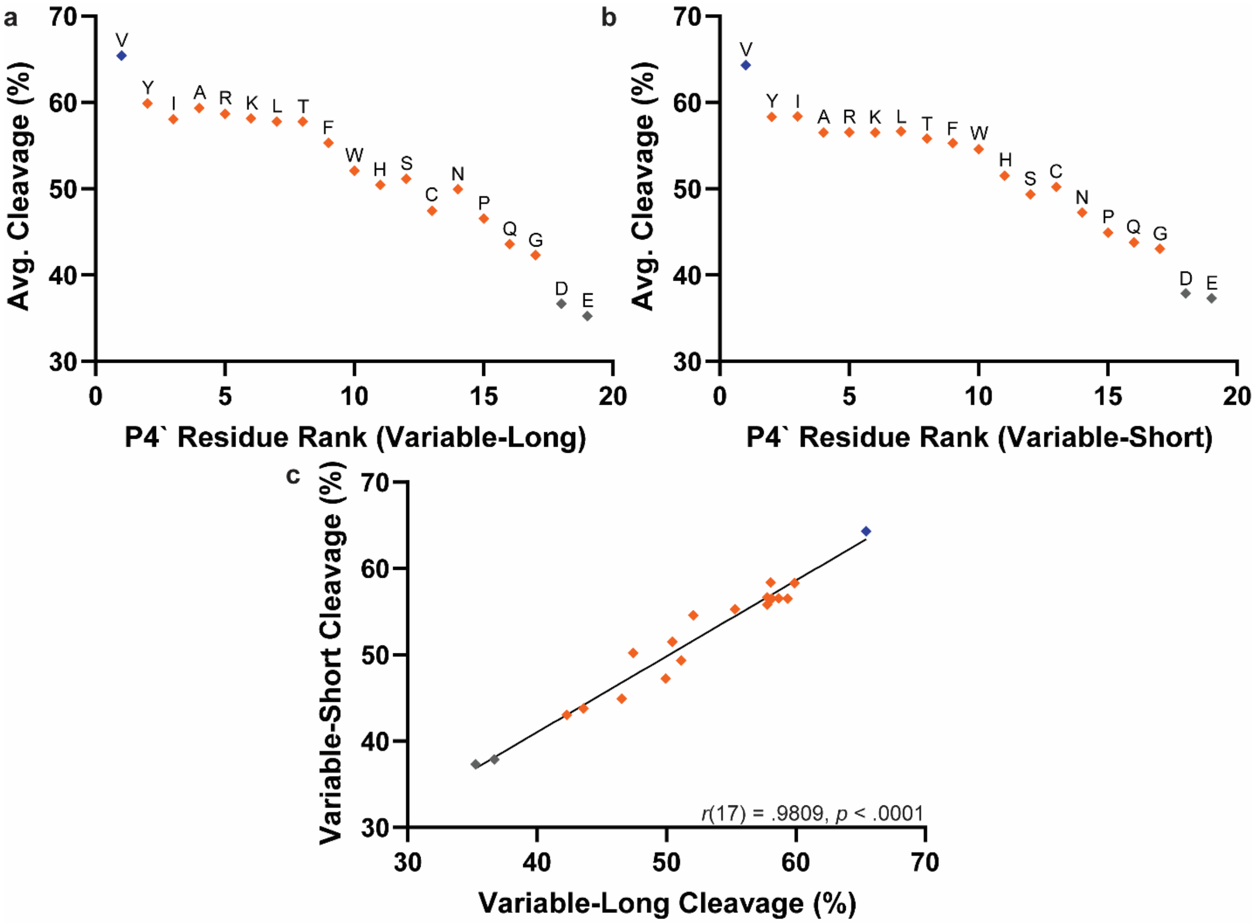

**Figure S8**: Estimated percent cleavage values from P4` subgroup analyses. (a-b) Percent cleavage values are presented for Variable-Long (a) and Variable-Short (b) libraries individually as a ranked list. (c) Variable-Long and Variable-Short library percent cleavage values are plotted against each other. The Pearson correlation coefficient is given, and the linear regression best-fit line is drawn to help visualize the linear relationship. Colors correspond to cleavage efficiency tiers as in **Figure 4** (blue = Tier 1; orange = Tier 2; gray = Tier 3).

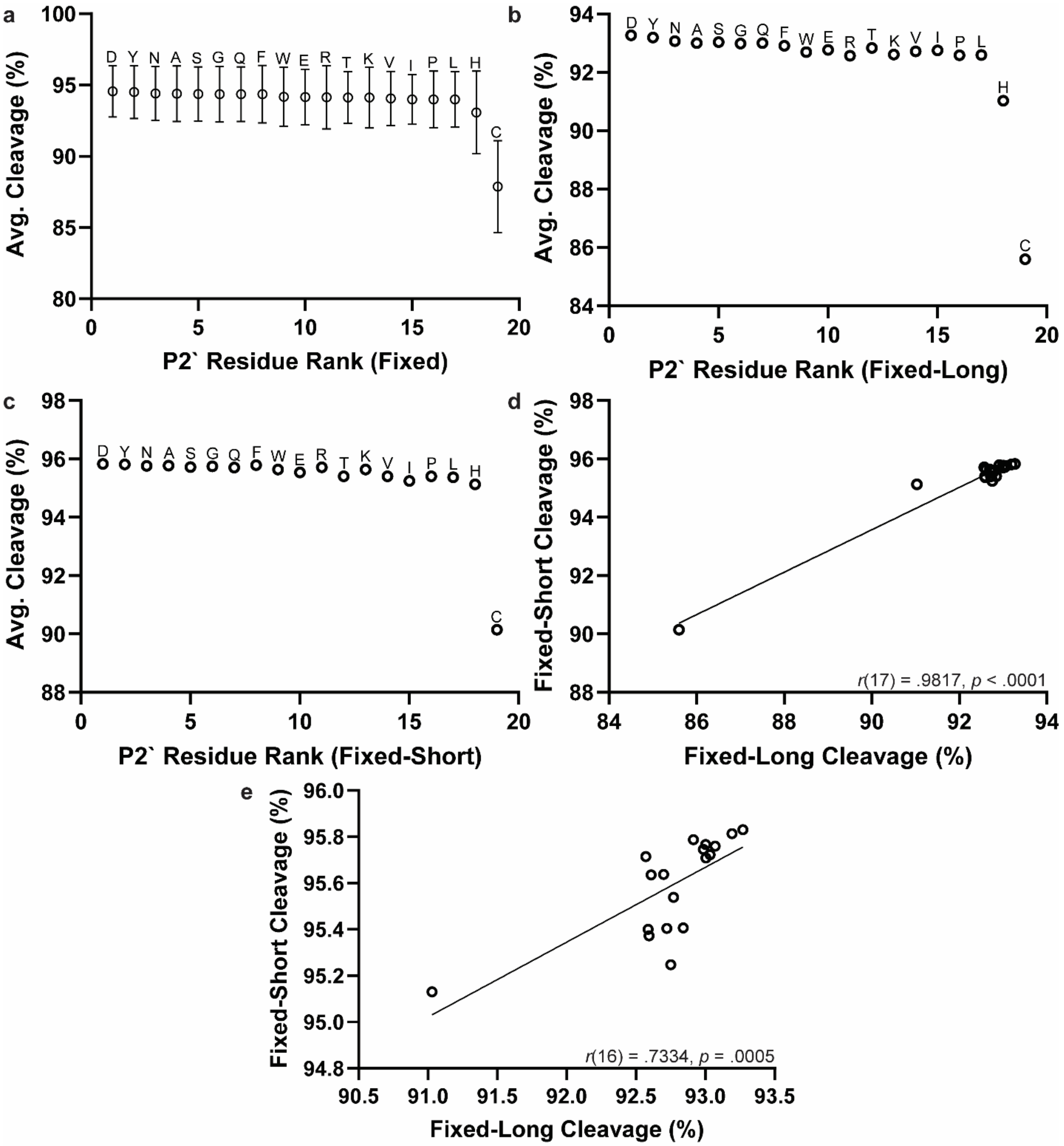

**Figure S9**: Estimated percent cleavage values from P2` subgroup analyses for Fixed libraries. (a-c) Percent cleavage values are presented as a ranked list for averaged values (a) or for Fixed-Long (b) and Fixed-Short (c) libraries individually. (d-e) Fixed-Long and Fixed-Short library percent cleavage values are plotted against each other either including (d) or excluding (e) Cys. The Pearson correlation coefficient is given, and the linear regression best-fit line is drawn to help visualize the linear relationship.

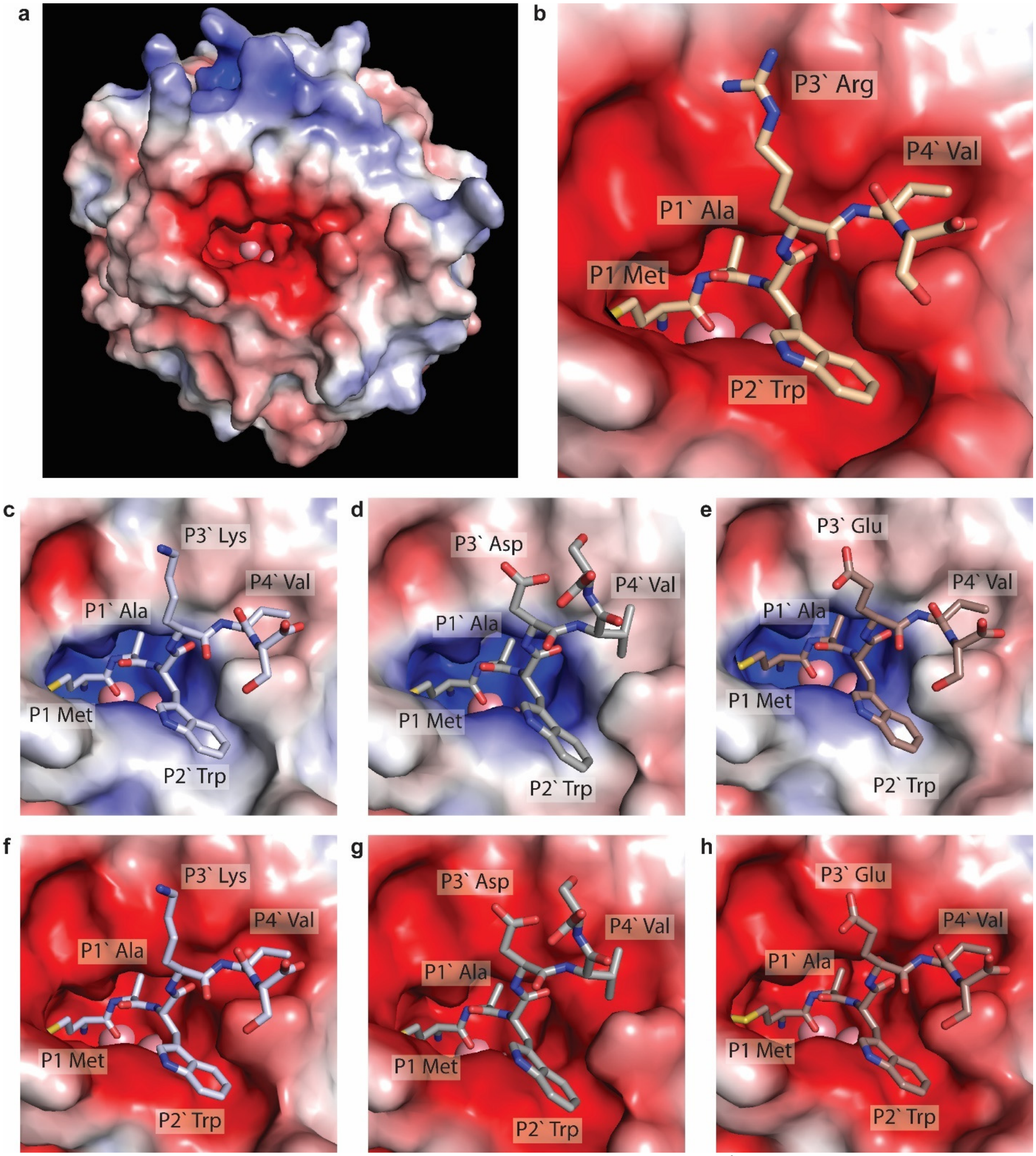

**Figure S10**: (a) Electrostatic potential surface of the MAP enzyme (PDB: 2MAT)[6] with binding pocket cobalt ions removed for electrostatic potential calculation (PyMOL 1.8 with APBS plugin).[7,8] (b) AlphaFold predicted structure described in **Figure 5** with cobalt ions removed for electrostatic potential calculation as in (a). (c-h) AlphaFold predicted structures of MAP (colored by electrostatic potential calculation either with (c-e) or without (f-h) cobalt ions included) in complex with short peptide MAWKVS (c,f) (ipTM = 0.96, pTM = 0.97), MAWDVS (d,g) (ipTM = 0.95, pTM = 0.96), or MAWEVS (e,h) (ipTM = 0.97, pTM = 0.97). Cobalt ions are displayed in all panels for visualization.

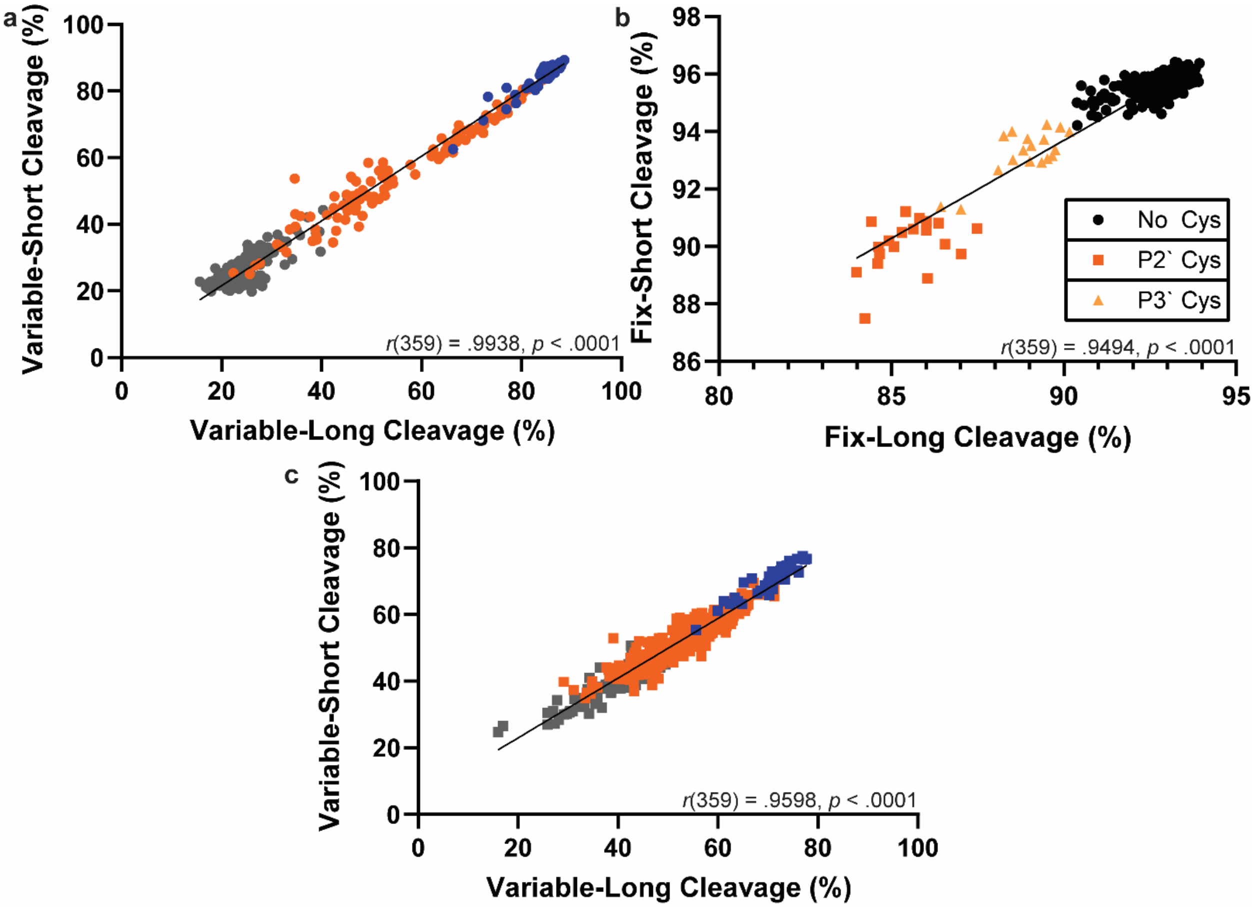

**Figure S11**: Estimated percent cleavage values with appropriate Long and Short libraries plotted against each other from the following two-residue combination subgroup analyses: (a) P1\` × P2\` Variable libraries, (b) P2\` × P3\` Fixed libraries, and (c) P2\` × P3\` Variable libraries (fixed P1\` Asn, Ile, Gln, Glu, Leu, or Asp). For each, the Pearson correlation coefficient is given, and the linear regression best-fit line is drawn to help visualize the linear relationship. (a+c) Colors correspond to cleavage efficiency tiers as in **Figure 4** (blue = Tier 1; orange = Tier 2; gray = Tier 3) for the P1\` (a) and P2\` (c) positions, respectively.

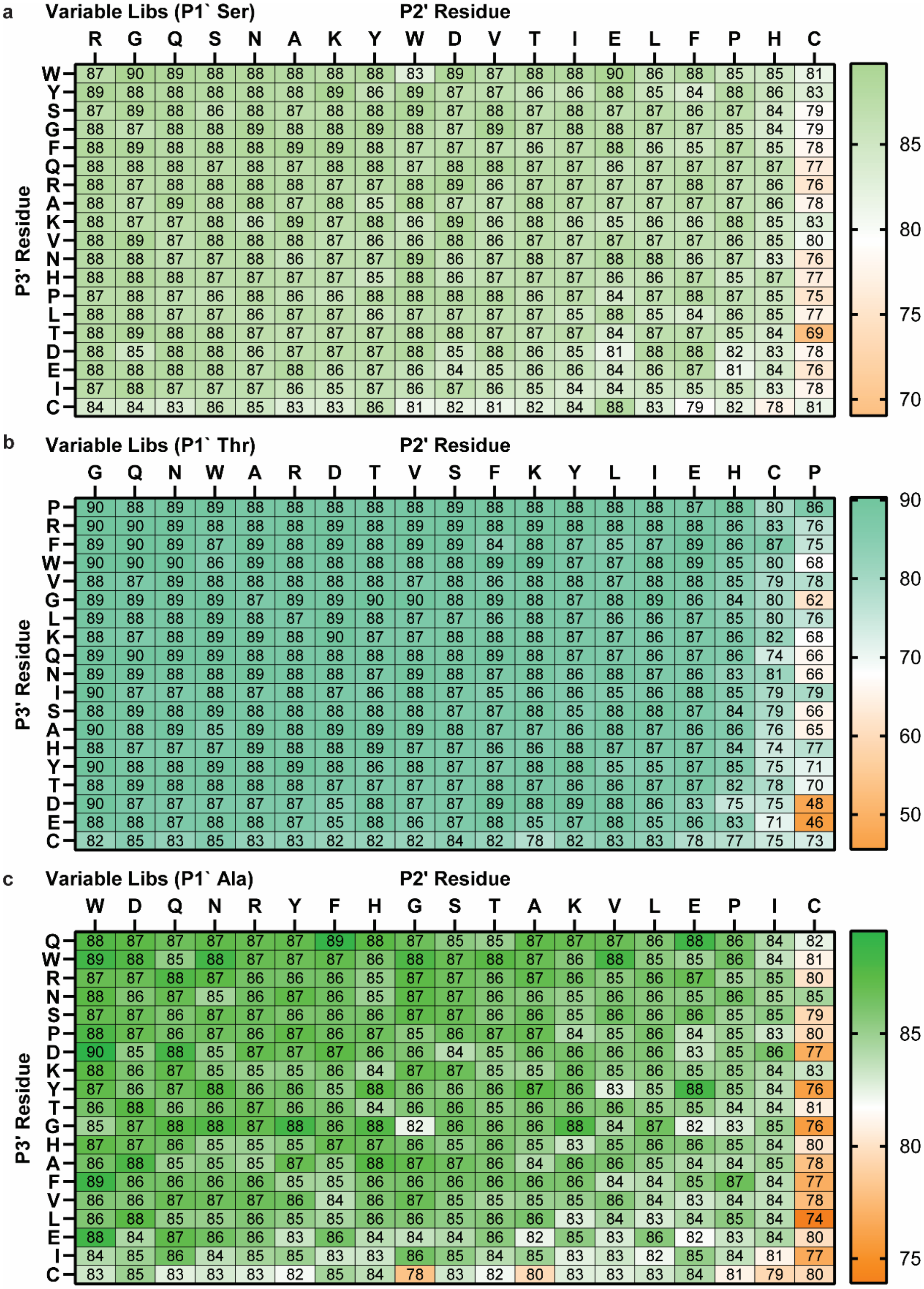

**Figure S12**: Heatmaps of percent cleavage calculated from subgroup analyses of two-residue combinations: P2` × P3` using averaged Variable libraries (fixed P1` Ser (a), Thr (b), or Ala (c)). Columns and rows are sorted by average values.

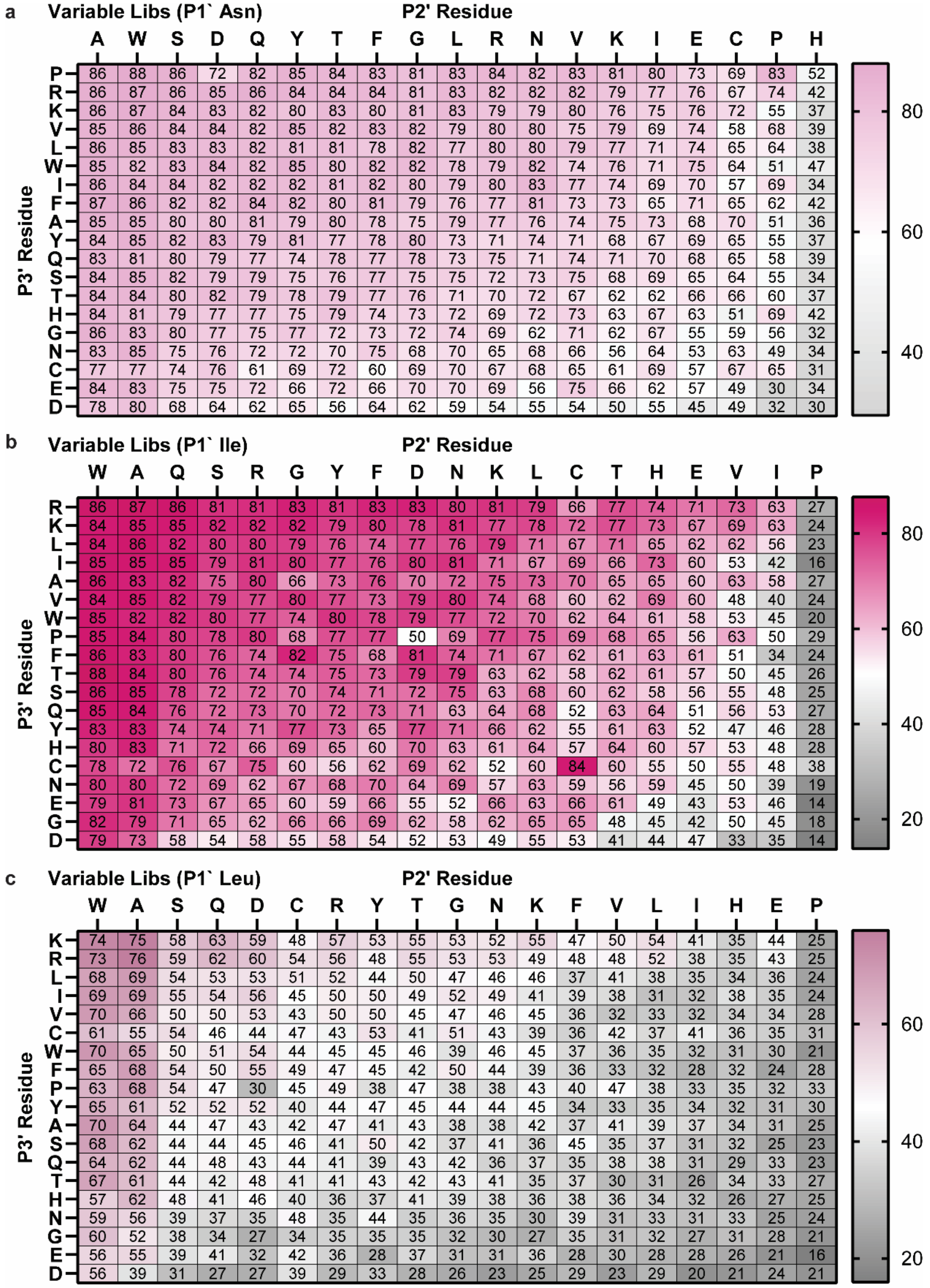

**Figure S13**: Heatmaps of percent cleavage calculated from subgroup analyses of two-residue combinations: P2` × P3` using averaged Variable libraries (fixed P1` Asn (a), Ile (b), or Leu (c)). Columns and rows are sorted by average values.

**Table S4**: Summary of previous literature investigating the substrate specificity of *E. coli* MAP. For each reference and substrate residue position, residues are placed in a blue (better cleavage), orange (intermediate cleavage) or gray (worse cleavage) category based on their influence on MAP cleavage as concluded by the authors. Unless specified, only *in vitro* assay results have been included.

| Ref | P1 | P1` | P2` | P3` | P4` |
|---|---|---|---|---|---|
| Ben-Basset *et al.*[9] | M | (A>G)/P(**1**) | M | | |
| | | I | G/S | | |
| | L/T/V/W/F/Y/I/R/S/A/E/G/P/ Mox/fM | [F/L/M/E/R/K](**2**) | P | | |
| Hirel *et al.*[10] | Only M tested | G/A>P/S/T/V>C | | | |
| | | N/D/L/I | | | |
| | | H/Q/E/F/M/K/Y/W/R | P | | |
| Frottin *et al.*[11] | M/Nle | A>S/Abu/G/P/C(**3**) | W/M/S | H/W | F/I |
| | | T/V/Nva | A/F/G/H/K/L/N/Q/R/V/Y/ Nva/Abu/D-Ser | G/K/M/N/R/S>A/F/I/L/Q/V/Y | K/S>G/M/N |
| | F/Mox/Nva/L>Abu/A | [I/N/D/M/L/F/Y/W/E/Q/K/R](**4**) | D/I/T>E>P | E/P/T>D | E |
| Merkel *et al.*[12] | M | G | | | |
| | Aha>Hpg | | | | |
| | | R | | | |
| Wang *et al.*[13](**5**) | | A>S | | | |
| | Aha>Hpg | G | | | |
| | | Q/E/H | | | |
| Wiltschi *et al.*[14] | M/Nle | A>G | | | |
| | Aha>Hpg | | | | |
| | Cpa | | | | |
| Xiao *et al.*[15] | Only M tested | A/S>G/Abu(**6**)/P(**7**) | E/D/N/Q/S/T/A/Abu/F/R(**8**) | Abu/S/T/V/A | |
| | | V/T | G/I/L/Nle/T/V/Y | | "No obvious selectivity" |
| | | D/E/F/H/I/K/L/N/Q/R/W/Y/Nle(**6**) | P/H(**9**)/W | [D/E](**10**)/G/H/W | |

**">"**: amino acid(s) on the left results in more efficient MAP cleavage than the amino acid(s) on the right

**Amino acid abbreviations**: Mox = Methionine sulfoxide, fM = N-formyl-Met, Nle = Norleucine, Nva = Norvaline, Abu = α-aminobutyrate, Aha = Azidohomoalanine, Hpg = Homopropargylglycine, Cpa = ß-cyclopropylalanine

(**1**) P1` Pro is not directly compared to either P1` Ala or P1` Gly
(**2**) Authors note that cleavage of some intermediate-sized side chain residues may be dependent on downstream sequence or tertiary structure
(**3**) Authors infer based on radius of gyration that Cys, which was incompatible with the main assay, will behave similar to Pro
(**4**) No peptides with P1` His appear to have been tested
(**5**) These results come from *in vivo* assays
(**6**) Since Cys and Met were not compatible with the main assay, Abu and Nle, respectively were used as surrogates
(**7**) Although P1` Pro performed poorly in the main assay, synthesized peptide was cleaved efficiently
(**8**) Although basic residues (Arg and Lys) were not compatible with the main assay, synthesized peptide with P2' Arg was cleaved efficiently
(**9**) Synthesized peptide performed better. Authors note P2' His can inhibit MAP activity at high peptide concentrations
(**10**) Authors note that having acidic residues in both the P2` and P3` positions was notably disadvantageous

### III. Kinetic Assay

#### A. Materials

For peptide synthesis, DMF (sequencing grade), EDT (ethane-1,2-dithiol), phenol, thioanisole, dimethyl sulfide, ammonium iodide, 1-hydroxybenzotriazole (HOBt), and DIC (N, N'-diisopropylcarbodiimide) were purchased from Sigma-Aldrich, Acros, Alfa-Aesar, TCI, or Fisher and used without further purification unless otherwise noted. All Fmoc-protected amino acids (except homopropargylglycine), piperidine, trifluoroacetic acid, and Oxyma® were purchased from Chem-Impex International, Inc. Fmoc-L-(homopropargyl)-Gly-OH was prepared in-house.[16] H-Rink Amide Chemmatrix® resin was purchased from Biotage and Sigma Aldrich. Milli-Q (MQ) ultrapure water and molecular biology-grade dimethyl sulfoxide were used for kinetic study.

#### B. Microwave-assisted Solid-Phase Peptide Synthesis

Automated solid-phase peptide synthesis (SPPS) was performed on the Liberty Blue peptide synthesizer (**Scheme S1**). The coupling and Fmoc deprotection were performed at 90 °C. Peptides were synthesized using DMF as a solvent. H-Rink amid Chem Matrix resin (0.42 mmol/g) was swelled in 10 mL of dry dichloromethane for 30 minutes before use in SPPS. Peptides were synthesized on a 0.05 mmol scale (120 mg resin), and each coupling was performed using a 0.2 M solution of Fmoc-amino acids (1.25 mL for each amino acid), 0.5 M Oxyma (0.5 mL), and 0.25 M DIC (1 mL) as coupling reagents. Fmoc-deprotection was done with 20% (v/v) piperidine in DMF containing 0.1 M Oxyma. The following amino acids were used in SPPS (all L-): Fmoc-Ala-OH (A), Fmoc-Asn(Trt)-OH (N), Fmoc-Glu-($^t$OBu)-OH (E), Fmoc-Gly-OH (G), Fmoc-Lys(Boc)-OH (K), Fmoc-Pro-OH (P), Fmoc-Ser($^t$Bu)-OH (S), Fmoc-Met-OH (M), Fmoc-Thr($^t$Bu)-OH (T), Fmoc-Tyr-($^t$Bu)-OH (Y), Fmoc-HPG-OH (X).

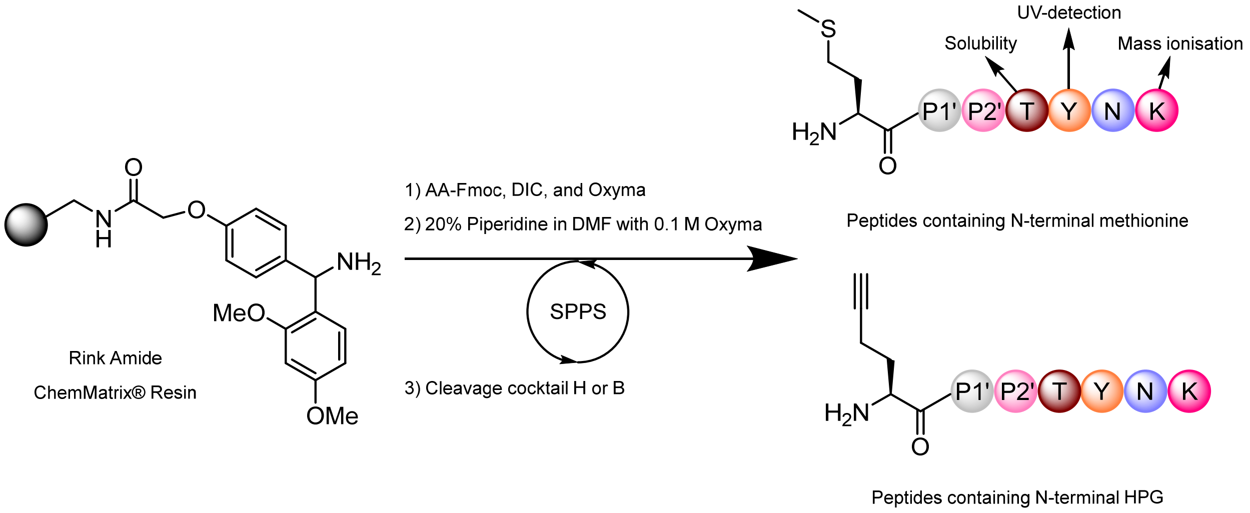

**Scheme S1**: SPPS overall process and design of sequence at positions 4-7.

#### C. Resin Cleavage and Purification

Following SPPS, resin was washed twice with 10 mL dichloromethane and transferred to a 50 mL conical tube. 10 mL of freshly prepared cocktail H (81% (v/v) trifluoroacetic acid, 5% (v/v) phenol, 5% (v/v) thioanisole, 2.5% (v/v) 1,2-ethanedithiol, 3% (v/v) water, 2% (v/v) dimethylsulfide, 1.5% (v/v) ammonium iodide) or cocktail B (88% (v/v) trifluoroacetic acid, 5% (v/v) phenol, 5% (v/v) water, 2% (v/v) triisopropylsilane) was added to the resin for Met-containing peptides or for HPG-containing peptides, respectively, and the tube was capped. After 1.5 hours at room temperature, the resin was filtered and washed twice with 5 mL of cleavage cocktail. The combined cleavage cocktail was evaporated under a stream of nitrogen and triturated with cold ether to obtain a crude pellet. The crude was dissolved in 4 mL of 50/50 water/ACN (containing 0.1% (v/v) formic acid), filtered through a 0.22-micron filter, and lyophilized. Peptides were then HPLC purified by a Waters XBridge OBD C18 column (10 x 250 mm) generally using a gradient of 1 to 10% (v/v) acetonitrile in water (containing 0.1% (v/v) formic acid) over 20 min (flowrate: 4 mL/min). After lyophilization, peptides were resuspended in 1250 µL water, quantitated by NanoDrop (**Table S5**), relyophilized, and resuspended in 27.8% (v/v) DMSO in water to a concentration of 5 mM. Purified peptides were checked by LC-MS using an Acquity UPLC BEH C18 column (2.1 x 150 mm, 1.7 µm, 130 Å) generally with 1 to 20% (v/v) acetonitrile in water (containing 0.07% (v/v) formic acid) over 7 minutes (flowrate: 0.3 mL/min).

**Table S5**: Yields of synthesized peptides after purification. `X` = HPG.

| Peptide sequence | Quantity (mg) |
|---|---|
| MATTYNK | 1.3 |
| MSWTYNK | 2.5 |
| MSETYNK | 5.5 |
| MAETYNK | 1.6 |
| MAWTYNK | 3.1 |
| MAPTYNK | 8.8 |
| MEPTYNK | 8.2 |
| MGPTYNK | 14.5 |
| MGTTYNK | 9.9 |
| XATTYNK | 2.2 |
| XSWTYNK | 4.1 |
| XSETYNK | 4.1 |
| XAETYNK | 5.5 |
| XAWTYNK | 3.7 |
| XAPTYNK | 9.4 |
| XEPTYNK | 6.8 |
| XGPTYNK | 9.6 |
| XGTTYNK | 8.6 |

**D. Linearity of MS-SIR Detection**

In preparation for kinetic assays, we first verified the linearity of peptide detection at assay concentrations using mass spectrometry in Single Ion Recording (SIR) mode. A representative peptide, XAETYNK (X = HPG), was prepared as a 20 mM stock solution, and 5 µL volumes of 2-fold serial dilutions were assayed by LC/MS down to 0.08 µM. Samples were run on the Waters Acquity/ZQ4000 single-quad MS, using an XBridge C18 BEH 1.7 um, 130 Å, 2.1 x 150 mm column in a gradient of 2-10% ACN in water (with 0.07 % formic acid) over 6 minutes, with MS in SIR mode with a cone voltage of 5 V. Linearity and good signal-to-noise were both observed in the range of 200 to 0.8 µM (**Figure S14-S15**).

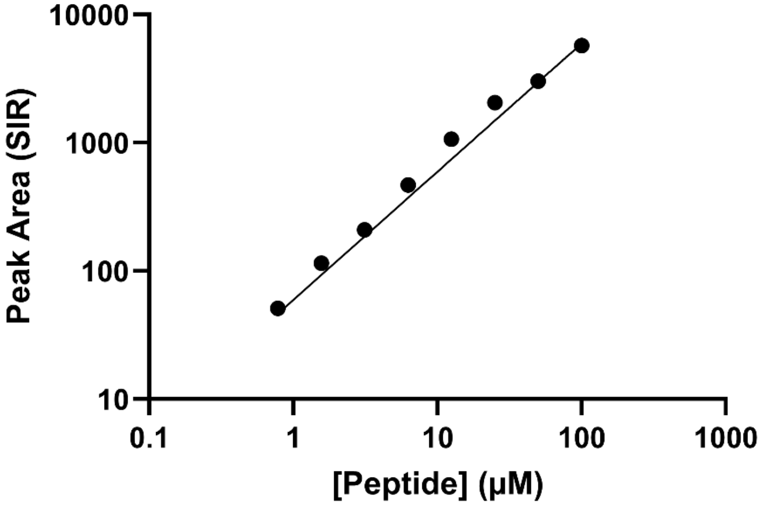

**Figure S14**: SIR peak area versus peptide concentration.

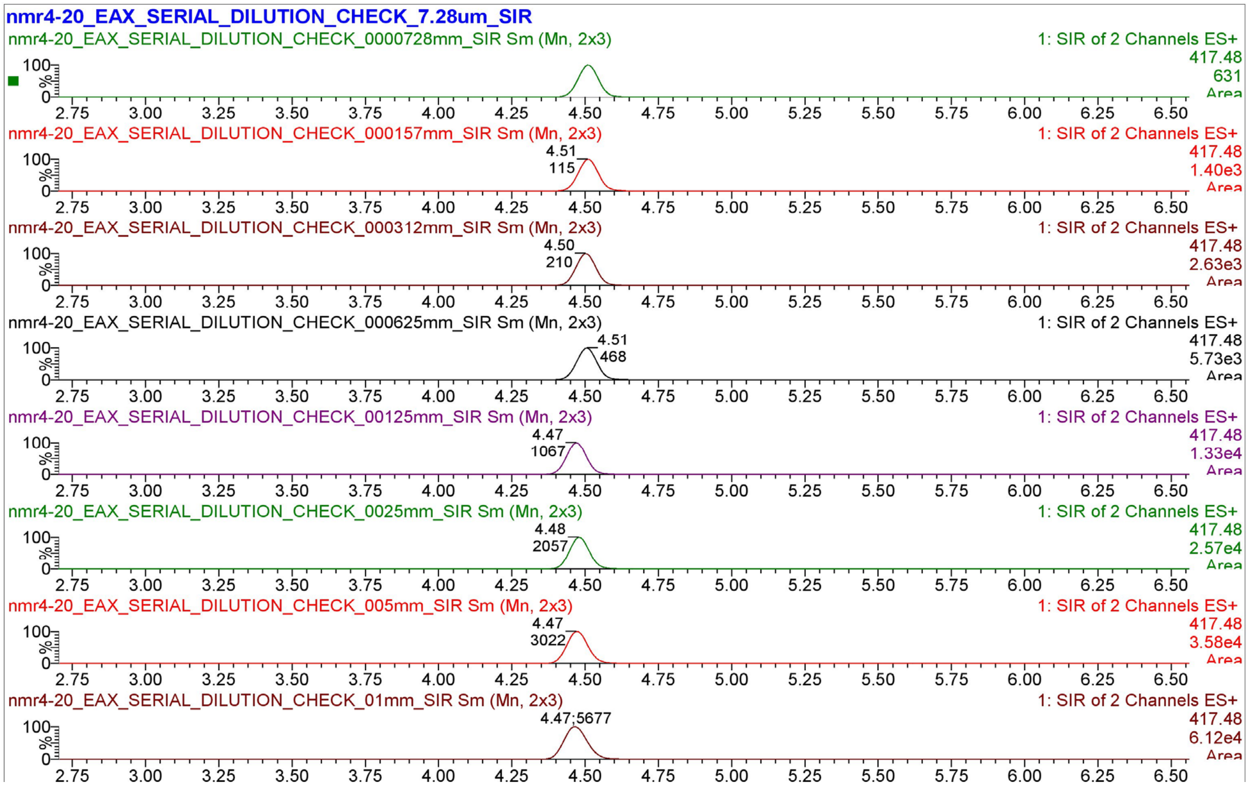

**Figure S15**: Representative integrated SIR chromatograms for linearity test.

### E. General Procedure for Peptide Cleavage Kinetics Assay

250 µL cleavage assay mixtures were prepared in triplicate at the following concentrations: 0.1 M phosphate buffer (pH 7.5), 0.2 mM $CoCl_2$, 0.43 µM MAP enzyme, and 0.18 mM peptide in 27.8% (v/v) DMSO in water for final 1% (v/v) DMSO in the assay mixture. The reaction mixture then stood at room temperature and aliquots were periodically removed. At each time point, a 20 µL aliquot was quenched with 5 µL 0.5 M EDTA (pH 8.0) and diluted with 1 mL water containing HOBt. 5 µL was injected into the LC-MS for quantitation of the uncleaved peptide, HOBt, and cleaved peptide by integration of SIR chromatograms at the corresponding retention time and m/z values. The HOBt at constant concentration in all samples served as a relative quantification standard to correct for fluctuations in the ~5 µL volume injected by the autosampler. Percent cleavage was calculated based on the decrease of [(peptide SIR peak area)/(HOBt SIR peak area)]. **Figure S16** shows a representative experiment, with gradients varying slightly depending on the peptide assayed.

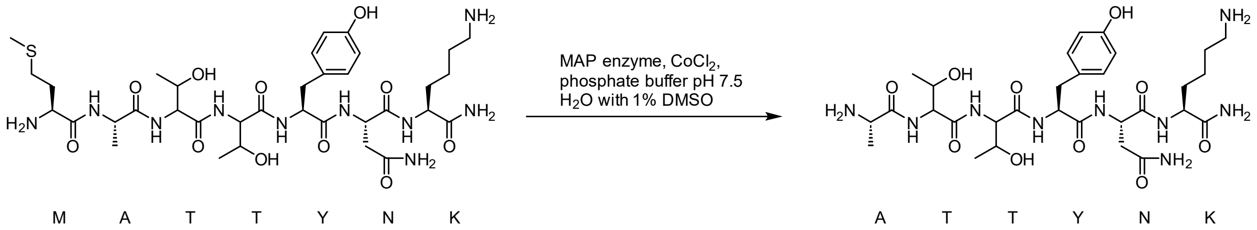

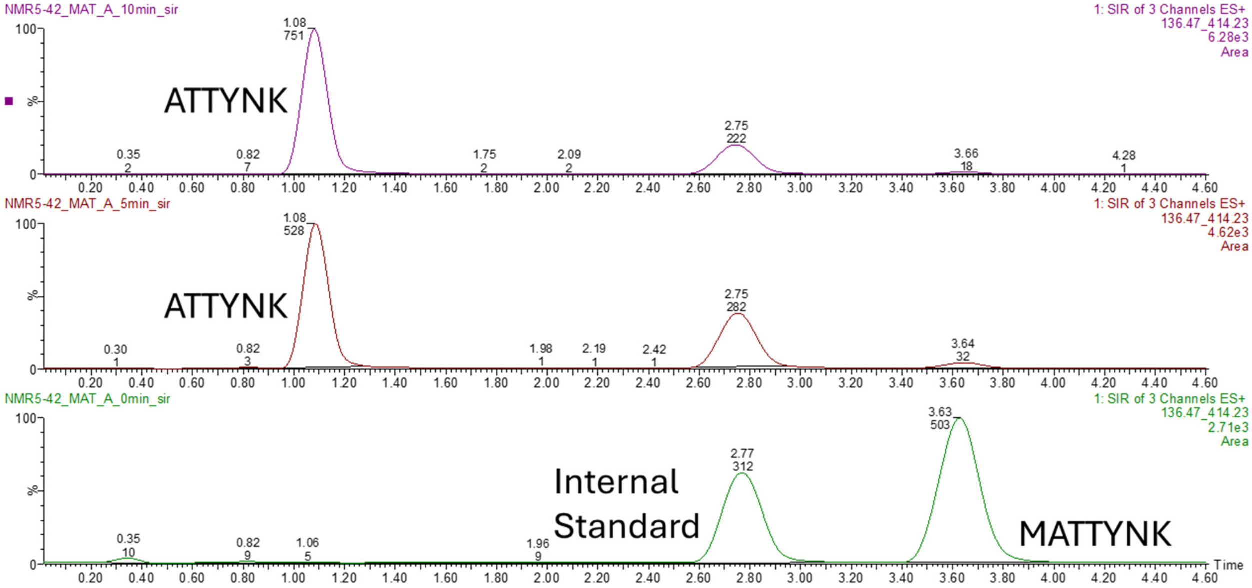

**Figure S16**: LC-MS profile for MATTYNK peptide cleavage experiment indicating different time points: 0 min, 5 min, and 10 min timepoints (from bottom to top).

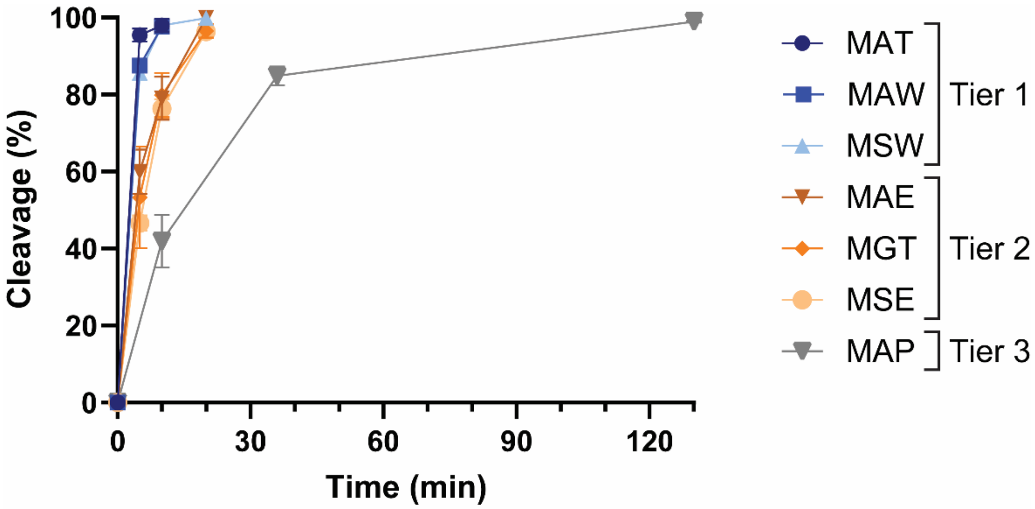

**Figure S17**: All data points of percent cleavage observed during time-course kinetic assay for synthetic heptapeptides containing N-terminal Met. All peptide sequences followed by -TYNK.

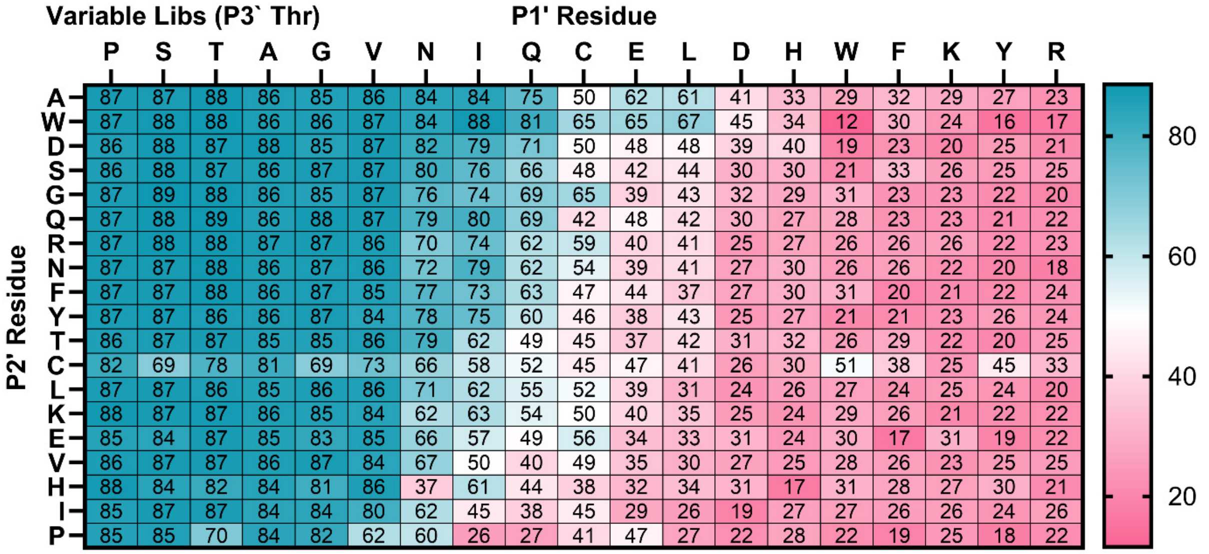

**Figure S18**: Heatmap of percent cleavage calculated from subgroup analysis of two-residue combination P1` × P2` using averaged Variable libraries (fixed P3` Thr). Columns and rows are sorted by average values.
